# Single-Cell Phylogenetic Tracing Maps the Landscape of Human Tumor Immune Cell Plasticity Driven by Microenvironment Factors-Mediated Epigenetic Reprogramming

**DOI:** 10.1101/2023.04.28.538645

**Authors:** Chuijin Wei, Liaoliao Dong, Shumin Xiong, Yunhan Tang, Ping Yu, Wenpei Liu, Lin Cheng

## Abstract

The tumor microenvironment harbors a complex ecosystem of immune cells, the heterogeneity of which has been extensively characterized, yet the developmental trajectories and plasticity of these populations remain poorly understood. Here, by integrating mitochondrial DNA mutations and chromosomal single-nucleotide polymorphisms inferred from single-cell RNA sequencing data, we established a robust lineage tracing framework to precisely reconstruct phylogenetic relationships among immune cells at single-cell resolution across multiple human cancers. This approach enabled us to systematically map the dynamics within the tumor immune landscape, revealing extensive plasticity including transitions between immune cell subtypes, transdifferentiation across lineages, and dedifferentiation toward progenitor states—all occurring with cancer-type specificity. Tumor microenvironment extrinsic factors were identified as key drivers of this plasticity, directing both the acquisition of transitional states and the orientation of fate changes. Mechanistically, these plasticity events were governed by profound epigenetic reprogramming, with global remodeling of chromatin accessibility and DNA methylation patterns underlying identity shifts. Our study provides a comprehensive atlas of immune cell plasticity in human tumors, establishes the molecular principles governing these dynamic processes, and reveals new opportunities for therapeutic intervention through manipulation of immune cell state or fate decisions.

## Introduction

Immune cells within the tumor microenvironment have been shown to play a crucial role in cancer development. Targeting of these cells as a part of immunotherapy against cancer has yielded significant success in recent years; however, several questions still remain unanswered. With the advent of single-cell analysis, a growing number of immune cell types and subtypes have been identified, contributing to our understanding of immune cell heterogeneity within tumors[1]. Different immune cell types exhibit varying functions as pro-inflammatory or anti-inflammatory agents during cancer progression[2]. Furthermore, different subtypes of the same immune cell type can exhibit contradictory roles in promoting or inhibiting cancer cell proliferation, invasion, and metastasis, as evidenced by the functions of type 1 and 2 macrophages[3], and type 1 and 2 neutrophils[4, 5].

The generation of heterogeneous immune cells has historically been ascribed to canonical hematopoiesis, a top-down hierarchical developmental process[6]. Once generated, immune cells typically acquire and maintain a specific function and phenotype, a widely accepted and well-known notion. However, the emergence of cell reprogramming techniques has challenged this paradigm[7]. Mature immune cells can now be transdifferentiated from one type to another or dedifferentiated into immature progenitors or stem cells *in vitro* by overexpressing exogenous transcription factors or through chemical compound treatment[8–12]. Under pathological conditions, a subset of immune cells can acquire plasticity, undergoing a transition from one subtype to another[13]. Within the cancer microenvironment, it has been reported that both natural killer (NK) cells and erythrocytes can transdifferentiate into myeloid-derived suppressor cells driven by cytokines[14, 15], and that B-cell precursors can transdifferentiate into macrophage-like cells[16]. Given the complexity of the microenvironment and the heterogeneity of immune cells in cancer, the question of whether other immune cells undergo dynamic state transitions or fate changes remains largely unanswered. A fundamental challenge is to construct a comprehensive map of these immune cell dynamics.

To decode cell dynamics and construct trajectories of cell state transitions and fate changes, lineage tracing based on exogenous barcodes or endogenous genetic codes is the gold standard protocol. The former method has been extensively employed in animal models and *in vitro* assays, while the latter method has been utilized in the analysis of human samples[17]. However, this latter method has focused on bulk cells and single endogenous genetic code, which has been found to be disadvantageous due to its limited resolution and accuracy. In this paper, we integrate two genetic tracing methods together: mitochondrial DNA (mtDNA) mutation and chromosome single nucleotide polymorphism (SNP) inferred from single-cell RNA sequencing (scRNA-seq) data. This approach enables the construction of phylogenetic trees of single immune cells in the tumor microenvironment across multiple cancers. Combining precise cell type definition inferred from the same single cell, we establish a comprehensive landscape mapping the dynamics of immune cells within the tumor microenvironment (Fig. 1A). Based on these maps, we report numerous immune cell state transitions and fate changes for the first time, and analyze the underlying mechanisms potentially manipulating these dynamics, including extrinsic factors and epigenetic reprogramming. This study not only provides the first comprehensive atlas of the precise dynamic trajectories of tumor microenvironment immune cells at single-cell resolution, but also enhances our understanding of the generation of cell heterogeneity in cancer and can potentially improve the efficiency of immunotherapy by manipulating the direction of immune cell changes.

**Fig. 1.**
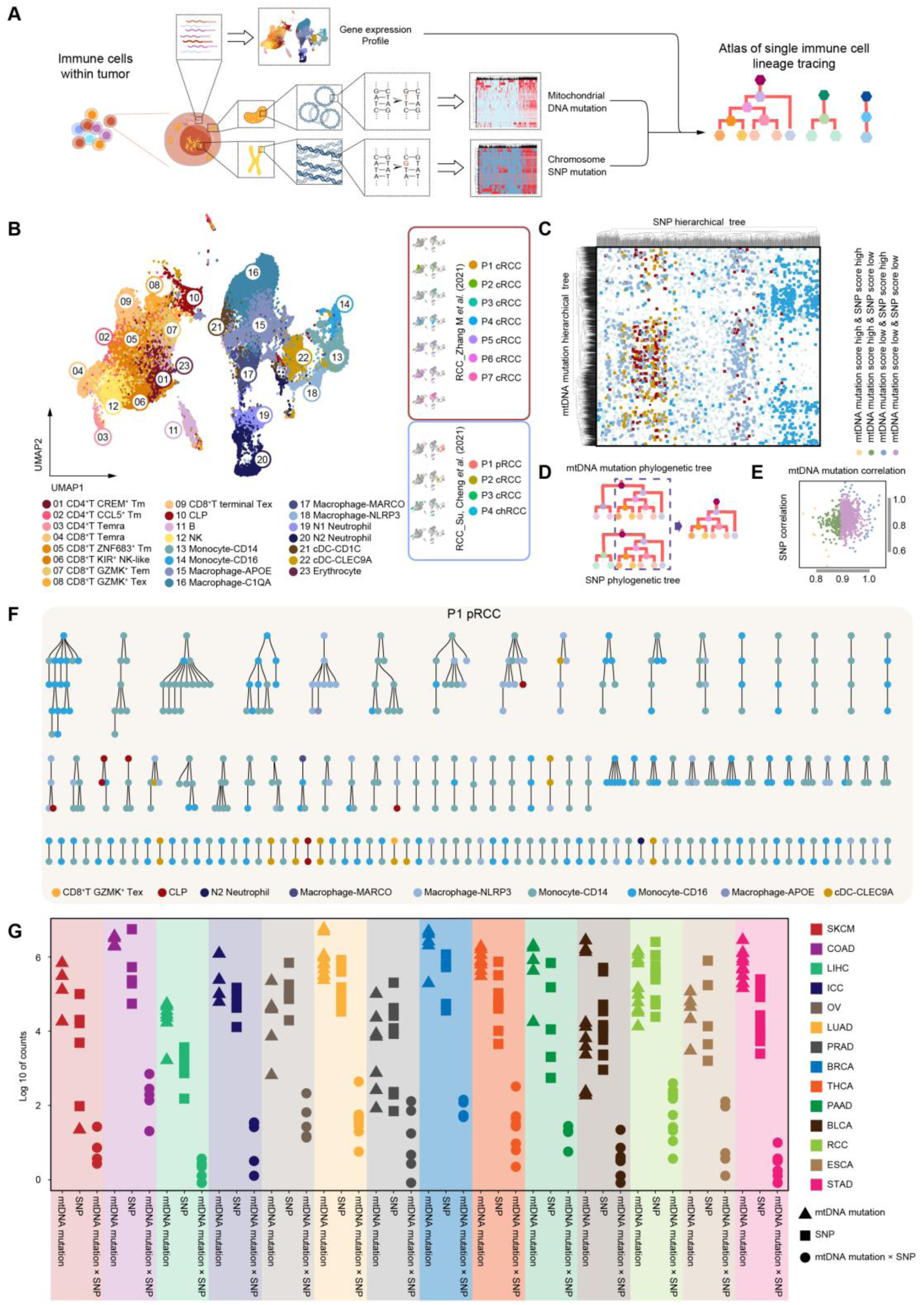
The lineage tracing method using transcriptome and genetic variation. (A) Schematic of the workflow for lineage tracing in pan-cancer using single-cell transcriptome, mitochondrial DNA mutation, and chromosome SNP mutation of immune cells. (B) UMAP plot showing the integration of two RCC scRNA-seq datasets (RCC_Zhang M *et al*., 2021 and RCC_Su, Cheng *et al*., 2021) on the left. UMAP plots displaying the distribution of each sample on the right. (C) Scatter plot displaying the immune cells of the P1 pRCC sample (RCC_Su, Cheng *et al*., 2021). Each dot represents a cell, with the x-axis indicating the rank of the SNP hierarchical tree and the y-axis indicating the rank of the mtDNA mutation hierarchical tree. A shorter distance between dots indicates a higher similarity in genetic information between cells. (D) Pattern diagram showcasing the phylogenetic tree (left) using mtDNA mutation and SNP, respectively, and the final phylogenetic tree (right) generated by overlapping the two methods. (E) Scatter plot illustrating the filtered two-cell pairs in the P1 pRCC sample (RCC_Su, Cheng *et al*., 2021). The filtration standard requires a mtDNA mutation correlation score of more than 0.8 and a SNP correlation score of more than 0.6. A high mtDNA mutation correlation score indicates a score above 0.9 but below 1.0, while a low mtDNA mutation correlation score indicates a score above 0.8 but below 0.9. A high SNP correlation score indicates a score above 0.8 but below 1.0, while a low SNP correlation score indicates a score above 0.6 but below 0.8. (F) Phylogenetic tree calculated by both mtDNA mutations and SNPs, depicting the process where ancestor cells transfer to descendant cells (from top to bottom) in the P1 pRCC sample (RCC_Su, Cheng *et al*., 2021). Each dot represents a cell and its color corresponds to the cell type. G. Scatter plot showing the number of different lineage tracing methods across various cancer types.

## Results

### Single Cell Immunophenotype Definition Paired with Combined Endogenous Genetic Codes to Construct Precise Phylogenetic Tree

Accurately defining cell types and subtypes is essential for further analysis. In scRNA-seq (10×) data from human renal cell carcinoma (RCC) samples, CD45^+^ cells were isolated and clustered into meta-clones and sub-clones. Employing UMAP analysis in conjunction with classical immune cell markers (Fig. S1A) enabled the identification of major immune cell types, including T cells, B cells, NK cells, monocytes, macrophages, neutrophils, dendritic cells (DCs), and common lymphoid progenitors (CLPs). Furthermore, major clusters were subdivided into more precise sub-clusters (Fig. 1B), which have been reported to exhibit diverse and specific functions[18–20]. RNA velocity analysis revealed that the primary developmental direction followed differentiation from progenitors into either lymphoid or myeloid cells[21]. However, subsequent analysis revealed that a significant proportion of immune cells did not align with this predominant trajectory. The velocity vectors for a subset of cells within each cluster deviated in multiple directions, with some even displaying opposing orientations (Fig. S1B). Additionally, pseudotime analysis revealed that numerous cells that clustered as the same cell type were not grouped together, but rather scattered in different branches defined as another cell type (Fig. S1C). Unsupervised clustering of mtDNA mutations and SNP of these cells, whether individually or collectively, further substantiated that diverse cell types could be closely or even jointly clustered (Fig. S1D and Fig. 1C), although the majority of the same cell types remained cohesive. These findings suggest that mature immune cells within the tumor microenvironment may exhibit plasticity, transitioning from one state to another or changing from one cell type to another at various frequency.

To gain a better understanding of the precise transitions or change directions between each immune cell pair, mtDNA mutations and SNP for each individual immune cell were inferred from scRNA-seq (10×) data[22, 23]. Subsequently, a phylogram was constructed, incorporating lineage alleles from each cell. In accordance with the protocol for tracing lineage, the offspring cell is expected to inherit the ancestral mutation points and accumulate further mutations. Applying this logic resulted in the isolation and verification of 4,351 or 5,047 cells in total 7,792 cells, respectively, within the ancestor-descendant hierarchy (Fig. S2A-J). In order to enhance the precision of the hierarchies, the two genetic trees were superimposed, and the cell pairs that exhibited a high degree of correlation in both phylogenetic trees were preserved and used for reconstruction (Fig. 1D, E). A precise ancestor-descendant single immune cell lineage tracing hierarchy was obtained, containing 596 cells in patient #1 RCC (P1 pRCC). Phylogenetic analysis revealed a series of notable transitions among different subtypes of macrophages, as well as instances of transdifferentiation between T cells and macrophages, and of dedifferentiation from macrophages to CLPs, in addition to cell expansion (Fig. 1F). Utilizing the refined lineage tracing protocol, which incorporates enhanced accuracy through effective false-positive filtering, we were also able to identify the paired cells across a total of 14 distinct cancer types (Fig. 1G).

It has been reported that diverse single cell sequencing methods can yield diverse insights concerning both cell type and genetic mutation. In this study, we applied our protocol to SMART-seq and scATAC-seq data and compared the cell type definition and phylogenetic tree among these two sources and scRNA-seq data[22]. We found that mtDNA coverage was comparable between SMART-seq and scATAC-seq, with both methods demonstrating superiority over scRNA-seq (Fig. S3A-D). Furthermore, the SNP counts from scATAC-seq, in the thousands, were higher than those from scRNA-seq, with SMART-seq, in the hundreds, having a lower count number (Fig. S3E-H). Upon combining the genetic codes for each data type, it was observed that the phylogenetic tree from SMART-seq contained a limited number of cell pairs, while the scATAC-seq tree exhibited several dozen cell pairs (Fig. S3I-L). However, the identification of specific cells was only possible in a limited number of cell pairs from SMART-seq, and the cells in the cell pairs from scATAC-seq exhibited ambiguity in cell type definition, despite the capacity to cluster them separately (Fig. S3M-T). Consequently, the subsequent phase of our study focused on comprehensive analysis of the scRNA-seq data.

### Validation of the Phylogenetic Tree Partially via T Cell Receptor Clonotypes

To validate our unexpected finding concerning T cell plasticity and dynamics, we conducted an in-depth analysis of the T cell receptor (TCR) in liver hepatocellular carcinoma (LIHC) [24]. A total of 3,309 TCR clonotypes were identified across 13 distinct immune cell types. Remarkably, we observed instances where different immune cell types and subtypes shared the same clonotype, strongly suggesting a shared genetic lineage (Fig. S4A, B). However, discerning the precise ancestor-descendant relationship proved challenging. Notably, 29 TCR clonotypes exhibited consistency with the phylogenetic tree derived from mtDNA mutations and SNP, displaying varying frequencies (Fig. S4C, D). Overall, our findings suggest that CD4^+^ T Tm cells, CD8^+^ GZMK^+^ Tex cells, CD8^+^ NK-like cells and CD8^+^ T Temra cells not only exhibited expansion but also underwent transitions to other cell subtypes or transdifferentiation into distinct immune cell types, although expansion events occurred at a higher frequency compared to state transitions and transdifferentiation processes (Fig. S4E).

For the transdifferentiation between CD8^+^ T Temra cell and neutrophil, we observed that two cells located in different clusters according to the UMAP plot acquired the exact same *TRA* and *TRB* gene sequences. This finding was supported by both mtDNA mutation analysis and SNP analysis, which further indicated a transdifferentiating direction from CD8^+^ T Temra cell to neutrophil (Fig. S4F-I). Similar results were observed for the transdifferentiation between CD8^+^ T Temra cell and B cell, with the former converting into the latter (Fig. S4K-N). Gene expression analysis of these cells revealed that the derived cells still retained partial key gene signatures of their original cell types (Fig. S4J, O). These data support our previously constructed phylogenetic tree based on scRNA-seq data-inferred two endogenous genetic codes. Although only a few immune cells exhibited the exact same genetic codes across all three methods, this is reasonable considering the probability of genetic code combinations (Fig. S4P).

### Landscape of Immune Cell Dynamics within Tumor Microenvironment

A comprehensive collection of scRNA-seq data from a variety of solid tumors was obtained, encompassing lung adenocarcinoma (LUAD), intrahepatic cholangiocarcinoma (ICC), skin cutaneous melanoma (SKCM), bladder urothelial carcinoma (BLCA), prostate adenocarcinoma (PRAD), thyroid carcinoma (THCA), esophageal carcinoma (ESCA), breast invasive carcinoma (BRCA), pancreatic adenocarcinoma (PAAD), stomach adenocarcinoma (STAD), colon adenocarcinoma (COAD), and ovarian serous cystadenocarcinoma (OV) (Fig. 2A). In total, there were approximately 0.5 million immune cells (Fig. S5A, B). Ten major cell clusters were identified, consisting of CD4^+^ T cells, CD8^+^ T cells, NK cells, B cells, monocytes, neutrophils, macrophages, mast cells, erythrocytes, and DCs. Furthermore, the majority of clusters were found to be further subdivided into multiple sub-clusters, a finding that aligns with prior reports (Fig. S5C).

**Fig. 2.**
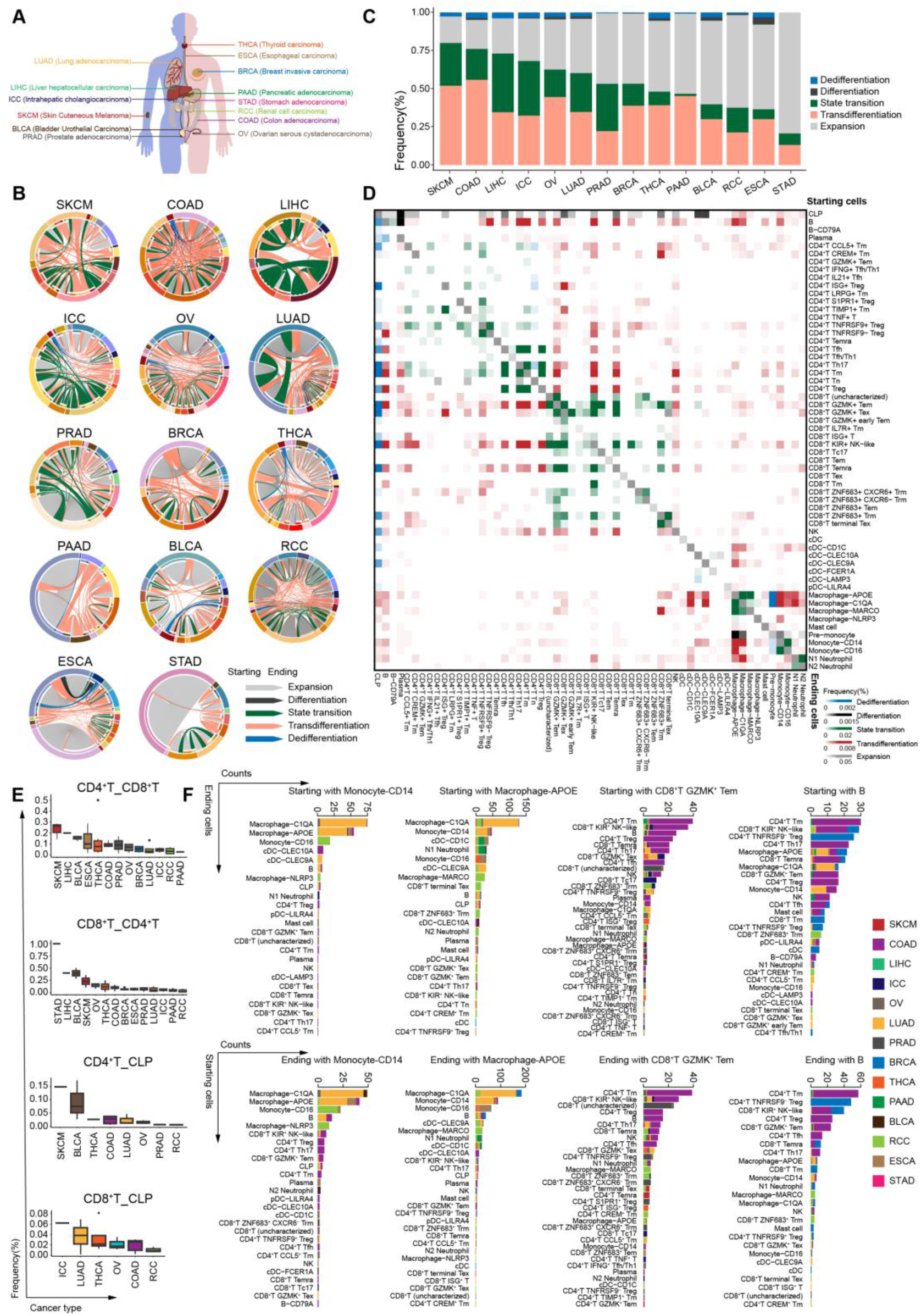
Dynamics of immune cells in pan-cancer at the single-cell resolution. (A) Pattern diagram showing cancer types involved in the pan-cancer analysis. The abbreviations are according to the TCGA database. (B) Circus plot showing the dynamics of the hematopoietic/immune cells in the 14 cancer types. Different colors of the arrows represent different cell change types. The arrows start with the starting cells and end with the ending cells. The colors of the outer ring represent the cell type of the starting cells, while the colors of the inner ring represent the cell type of the ending cells. The colors of cell type correspond to the cell type in Fig. S5. (C) Bar plot illustrating the frequency of the five cell change types among each cancer type. (D) Heatmap showing the frequency of cell change types in the pan-cancer analysis. The cell types in the rows are the starting cells, and the cell types in the columns are the ending cells. Different colors represent different cell change types. The frequency is calculated through dividing the counts of each cell change type by the total number of all cell change types. (E) Box plots showing the proportions of the four cell change types in major lineages across various cancer types. (F) Bar plots reporting the counts of starting cells and ending cells that changed from the labelled cell types across various cancer types.

We generated circus plots that represented every single immune cell type and their lineage relationships (Fig. 2B). Taking RCC as an example, the plots exhibited a clear indication that cell expansion constituted more than half of the observed relationships across the majority of cell types, particularly those comprising relatively higher cell numbers. State transitions between different cell subtypes within the same cell type were also observed. Surprisingly, cell transdifferentiation between different cell types occurred at a higher frequency compared to cell differentiation. Both state transitions and transdifferentiations were unidirectional or bidirectional. Additionally, a few mature immune cells could dedifferentiate into progenitors, although at a low frequency. All of the above findings illustrate that the function and phenotype of immune cells within RCC are dynamic, and the single-cell endogenous genetic mutations can help to decisively depict the lineage tracing direction of these dynamics.

### Statistics and Characterization of Tumor Microenvironment Immune Cell Dynamics

According to the individual dynamic landscapes of immune cell changes generated for each of the tumors, the frequencies of immune cell transdifferentiation, state transition, and dedifferentiation were significantly higher in SKCM, COAD, LIHC, ICC, OV and LUAD compared to other tumors, which had a higher frequency of immune cell expansion (Fig. 2C). The low frequency of immune cell differentiation and dedifferentiation could be ascribed to the restricted presence of hematopoietic stem or progenitor cells, including a small number of CLPs, within these tumors.

Immune cell expansion was observed across almost all immune cell types and subtypes, while differentiation was a well-known phenomenon. Consequently, the remaining three immune cell state transitions and fate changes were subjected to more exhaustive analysis (Fig. 2D). The analysis revealed that state transitions among CD4^+^ T cell subtypes, CD8^+^ T cell subtypes, macrophage subtypes, neutrophil subtypes, and monocyte subtypes occurred at a high frequency. Similarly, the frequencies of transdifferentiation between CD4^+^ T cells and CD8^+^ T cells, as well as between macrophages and neutrophils, were also high (Fig. 2E and Fig. S6A). Generally, both cell fate changes occurred at a higher frequency within lymphoid cells or myeloid cells. For example, transdifferentiation from CD4^+^ T cells to CD8^+^ T cells and vice versa was predominantly observed in SKCM, LIHC and BLCA, with a frequency of more than 10%. Transdifferentiations between macrophages and monocytes, macrophages and neutrophils, were mainly observed in LUAD, RCC and PAAD. However, transdifferentiation between these two major cell clusters, lymphocyte and myelocyte, occurred at a low frequency and was confined to minor cell subtypes, including CD4^+^ T Tm cells and monocytes, B cells and macrophages.

Interestingly, conventional DCs exhibited minimal transitions among their subtypes and rarely transdifferentiated from or to other immune cell types, with the exception of macrophages and monocytes. By contrast, B cells exhibited higher cell fate change activity. Bidirectional transdifferentiation between B cells and CD4^+^ T Tm cells, CD8^+^ T KIR^+^ NK-like cells, CD4^+^ T Treg, CD4^+^ T Th17, and CD8^+^ T GZMK^+^ Tm cells were mainly observed in COAD. The transdifferentiation between B cells and CD4^+^ T TNFRSF9^-^ Treg cells occurred mainly in BRCA (Fig. 2F). In LUAD, the predominant cell fate changes occur within myeloid cells, including monocytes and macrophages. Neutrophil transdifferentiation primarily occurred from or into macrophages in RCC, PRAD, and OV (Fig. S6B).

### Immune Cells Acquire Plasticity through Intermediate Cell State

Typically, cells undergoing a state transition or fate change require a degree of plasticity before the transformation can occur[25]. In order to ascertain whether immune cells exhibit this degree of plasticity, a comparative analysis of transcriptional signatures was conducted between lymphoid and myeloid cells. A thorough examination of the discrete values for each cell type and subtype revealed that specific lymphoid cells, including B cells from BRCA and THCA, CD4^+^ T CCL5^+^ Tm cells from OV, and CD8^+^ T Temra cells from PRAD, exhibited a partial acquisition of myeloid cell signatures. In contrast, a significant proportion of subtypes, such as macrophage-C1TA from LIHC, macrophage-APOE from PRAD, and monocyte-CD14 from ICC exhibited acquired lymphoid cell signatures (Fig. 3A). The presence of these mixed transcriptional signatures indicates the existence of intermediate cells that have acquired plasticity. The highest discrete value was observed in CLPs, while most DCs had the lowest value. These findings are consistent with the cell fate change pattern depicted in Fig. 2D.

**Fig. 3.**
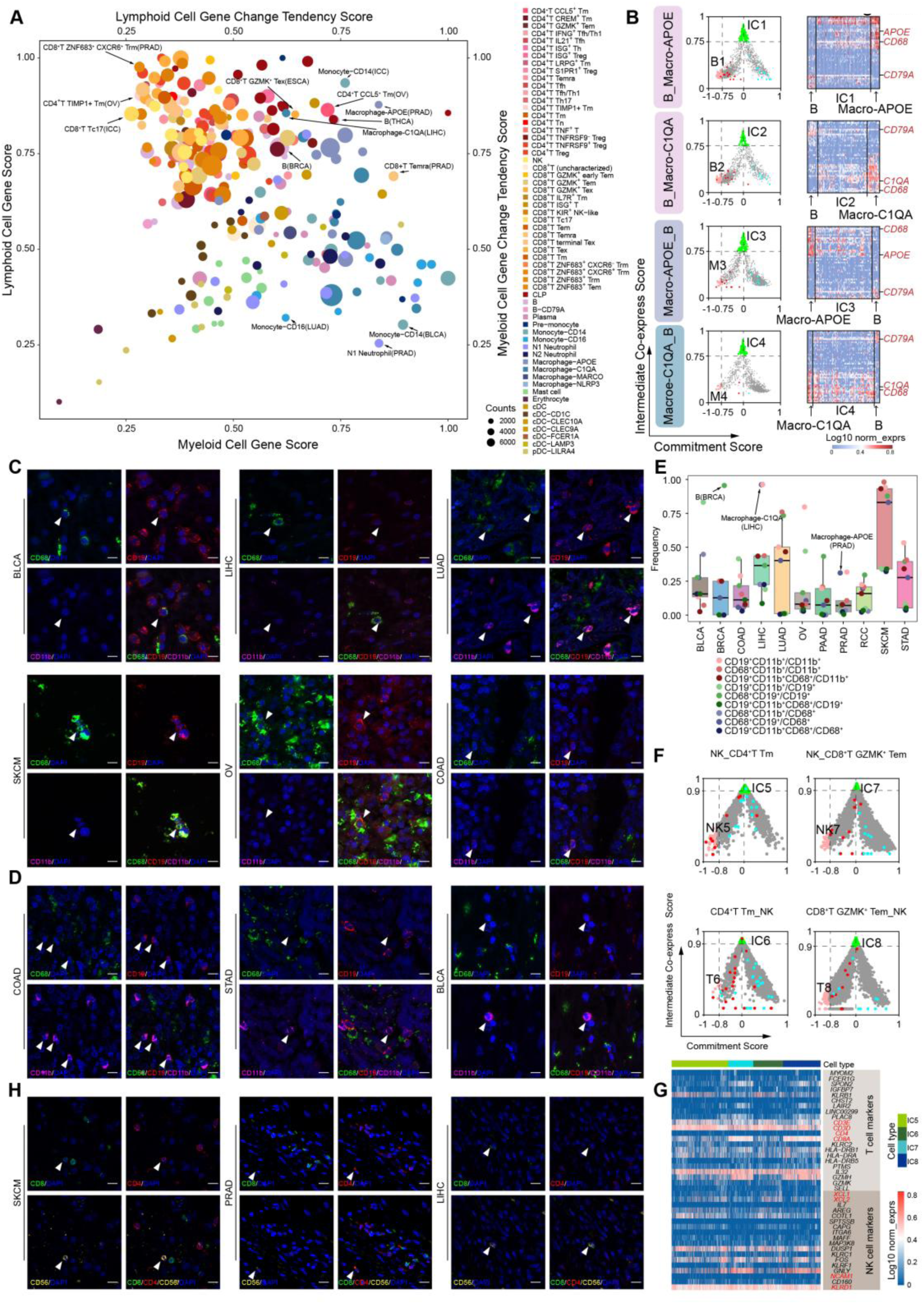
Intermediate state of immune cell plasticity. (A) Scatter plot illustrating the gene expression of the hematopoietic/immune cells and their potential ability to change. Each dot represents one cell type of the 14 cancer types and its size is proportional to the number of cells in that particular cell type. For myeloid cell types, the x-axis represents myeloid cell gene score, indicating their gene expression, and the y-axis represents myeloid cell gene change tendency score, indicating whether these cells tend to change. For lymphoid cell types, the y-axis represents lymphoid cell gene score, indicating their gene expression, and the x-axis represents lymphoid cell gene change tendency score, indicating whether these cells tend to change. Dendritic cells’ x-axis represents lymphoid cell gene change tendency score, while the y-axis represents myeloid cell gene change tendency score. Erythrocytes’ x-axis represents myeloid cell gene score, while the y-axis represents lymphoid cell gene score. (B) Scatter plot showing the change process of transdifferentiation between B and macrophage-APOE or macrophage-C1QA. The abscissa represents B/macrophage commitment score and the ordinate represents Intermediate Co-express Score. Cells with an x-value less than -0.75 are defined as potential starting cells, and cells with a y-value greater than 0.75 are defined as ICs. Green dots represent intermediate cells and pink dots represent potential starting cells. Red dots and blue dots represent the starting and ending cells of the transdifferentiation which we discovered in pan-cancer using the method we built in this study, respectively. The heatmaps on the right show the specific genes of B cells, macrophages and the intermediate cells. (C) Representative examples stained by multiplex immunofluorescence to show the co-expression of CD68 and CD19 (white arrows). The scale bar represents 20 μm. (D) Representative examples stained by multiplex immunofluorescence to show the co-expression of CD11b and CD19 (white arrows). The scale bar represents 20 μm. (E) Box plots showing the proportions of double-positive cells across various cancer types. (F) Scatter plot showing the change process of transdifferentiation between NK and CD4^+^ T Tm or CD8^+^ T GZMK^+^ Tem. The abscissa represents NK/T cell commitment score and the ordinate represents Intermediate Co-express Score. Cells with an x-value less than -0.8 are defined as potential starting cells, and cells with a y-value greater than 0.9 are defined as intermediate cells. Green dots represent intermediate cells and pink dots represent potential starting cells. Red dots and blue dots represent the starting and ending cells of the transdifferentiation which we discovered in pan-cancer using the method we built in this study, respectively. (G) Heatmap showing the NK cell markers and the T cell markers of the intermediate cells. (H) Representative examples stained by multiplex immunofluorescence to show the co-expression of CD56 and CD4 or CD8 (white arrows). The scale bar represents 20 μm.

To further validate the generation and characteristics of plastic cells, we calculated the intermediate co-expression score of the starting and ending cells based on the cell change pairs identified earlier[26]. Specifically focusing on the cell changes between B cells and macrophages, we discovered intermediate cells, designated as intermediate cell (IC) 1 and IC2, which transdifferentiate from B cells to macrophage-APOE and macrophage-C1QA, respectively. Conversely, intermediate cells designated as IC3 and IC4 were identified in the reverse direction, from individual macrophages to B cells (Fig. 3B). The IC1s, IC2s, IC3s and IC4s all co-expressed B cell-specific gene *CD79A* and macrophage-specific gene *CD68*, *APOE* or *C1QA*. To further investigate the intermediate cells within the tumor microenvironment, we performed multiplex immunofluorescent analysis, which confirmed their presence (Fig. 3C). Our data revealed that these intermediate cells co-express the B cell-specific marker CD19 along with the macrophage-specific marker CD68 in BLCA, LIHC, LUAD, SKCM, OV, and COAD. Furthermore, in COAD, STAD, and BLCA, we also identified a population co-expressing CD19 and the neutrophil marker CD11b, revealing not only the presence but also the phenotypic diversity of these intermediate cells (Fig. 3D). Statistically, the results revealed a greater prevalence of intermediate cells in SKCM, LUAD, and LIHC, as evidenced by multiplex immunofluorescence (Fig. 3E). It is noteworthy that the high cell change tendency score for B cells in BRCA, macrophage-C1QA in LIHC and macrophage-APOE in PRAD (Fig. 3A) was further corroborated by a high frequency of intermediate cells observed via multiplex immunofluorescence.

We extended our plasticity analysis to neutrophils and macrophages/monocytes. Both cell types exhibited minimal transdifferentiation capacity, as quantified by low cell change tendency scores (Fig. S7B). Correspondingly, we identified the intermediate cells from the cell changes among macrophage subtypes and neutrophil subtypes (Fig. S7C). Corroborating this finding, only a limited number of intermediate cells co-expressing the macrophage-specific marker CD68 and the neutrophil-specific marker CD11b were detected in PRAD and OV (Fig. S7D). Furthermore, the ratio of CD68^+^ CD11b^+^ double-positive cells to total CD68^+^ macrophages was low across most cancer types (Fig. 3E). Together, these data from both scoring and imaging analyses demonstrate a low transdifferentiation potential between neutrophil and macrophage lineages.

For cell change between CD4^+^ T cells and CD8^+^ T cells, some CD4^+^ T cell subtypes acquire CD8^+^ T cell characteristics and vice versa (Fig. S7A). Similarly, for transdifferentiation between CD4^+^ T Tm cells and NK cells, and between CD8^+^ T GZMK^+^ Tem cells and NK cells, ICs are present in all transdifferentiating pairs and directions (Fig. 3F). Intriguingly, the gene expression pattern of these ICs closely resembles that of natural killer T (NKT) cells[27], which express both NK cell markers (*KLRD1, KLRF1, XCL2*) and T cell markers (*CD3D, CD3E, IL32*) (Fig. 3G). We also identified intermediate cells co-expressing the NK cell-specific marker CD56 together with the T cell-specific markers CD4 or CD8 via multiplex immunofluorescence in SKCM, PRAD, and LIHC (Fig. 3H). The high frequency of these dual-positive cells suggests robust transdifferentiation potency between NK and T cell lineages, a finding corroborated by high plasticity scores (Fig. S7I). Moreover, we also found the intermediate cells in the state transition from CD4^+^ T Th17 cells into CD4^+^ T Treg cells (Fig. S7E) and the transdifferentiation between CD4^+^ T cells and CD8^+^ T cells (Fig. S7F). In line with the high plasticity observed in B cells, we detected numerous intermediate cells co-expressing the B cell-specific marker CD19 with the T cell-specific markers CD4 or CD8 (Fig. S7G, H, I), demonstrating active transdifferentiation between B and T cell lineages.

Taken together, our multimodal analysis confirms the pervasive and dynamic nature of immune cell plasticity within the tumor microenvironment. We identified intermediate cells across a spectrum of cell changes, including the transdifferentiation of B cells and macrophages, T cells and NK cells, and state transition among T cell subsets and myeloid lineages. These findings collectively establish cell plasticity as a fundamental feature of the tumor immune ecosystem.

### Tumor Microenvironment Extrinsic Factors Contributing to Immune Cell Plasticity

To determine whether tumor microenvironment extrinsic factors played a role in modulating immune cell fate changes, we made a ligand-receptor analysis which revealed that these changing cells selectively interacted with microenvironmental cells, including malignant epithelial cells, fibroblasts, endothelial cells, and other immune cells, and were influenced by the soluble factors secreted by these cells. We compared the intermediate cells and the starting cells of the cell changes to the cells that underwent expansion to identify the most potential factors contributing to the cell plasticity. This analysis identified several key factors potentially driving specific lineage transitions. The top factors potentially influencing the cell changes among macrophage subtypes and neutrophil subtypes were APOE and CSF1, along with FGF2, TGFβ1, INFγ, TNF, and CCL3. The cells that underwent fate changes among B cell and macrophage subtypes were potentially influenced by TGFβ1, AGT, CCL3, INFγ, and TNF. Moreover, our data confirms previous report that TGFβ1 induces the state transition from CD4^+^ T Th17 cells into CD4^+^ T Treg cells[28] (Fig. S8A). Furthermore, a combination of factors, including TGFβ1, FGF2, and JAG1, appeared to be associated with the generation of NKT ICs from various cell types[29] (Fig. 4A).

**Fig. 4.**
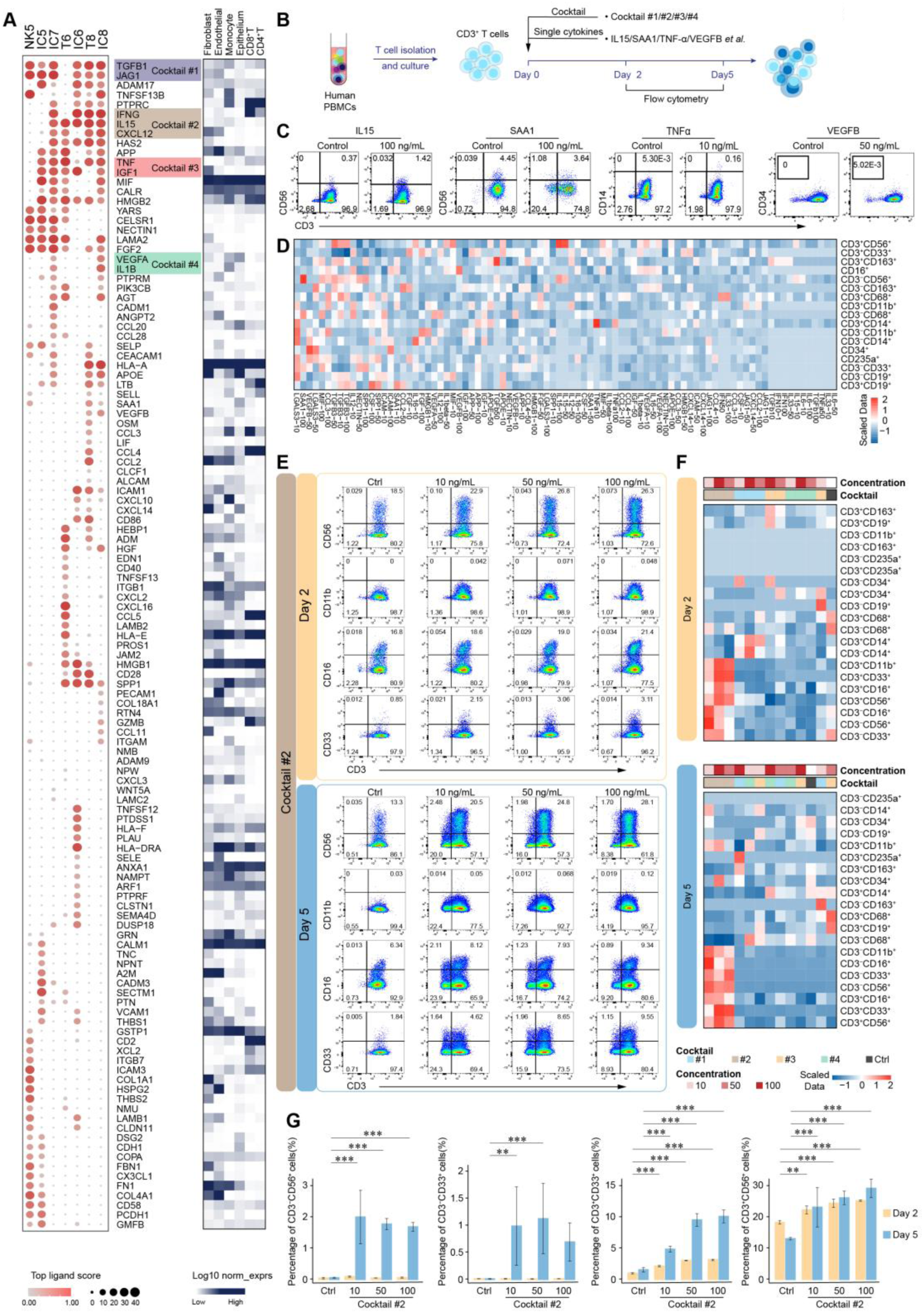
Tumor microenvironment factors contributed to immune cell plasticity. (A) Bubble plot illustrating the possible ligands driving the potential starting cells and intermediate cells in Fig. 3F against expansion NK cells and expansion T cells respectively, sent by other immune cells and parenchymal cells. The top 40 ligands are listed for each cell type and the heatmap shows the expression of ligands in the immune cells and parenchymal cells. NK7 is not listed because it has no differential gene with the expansion NK cells. (B) Experimental diagram of human primary T cell stimulation with individual cytokines and cytokine cocktails in (A). Flow cytometry was performed on day 2 and day 5. (C) Representative flow cytometry plots showing phenotype of T cells treated with individual cytokines on day 5. CD3^+^ CD56^+^ cells increased with IL15 treatment at 100 ng/mL. CD3^-^ CD56^+^ cells increased with SAA1 treatment at 100ng/mL. CD3^+^ CD14^+^ cells increased with TNFα treatment at 10 ng/mL. CD34^+^ cells increased with VEGFβ treatment at 50 ng/mL. (D) Heatmap showing distinct T cell phenotypes induced by individual cytokines at different concentrations on day 5. (E) Representative flow cytometry plots showing phenotype of T cells treated with cocktail #2 at different concentrations on day 2 and day 5. (F) Heatmap showing distinct T cell phenotypes induced by cytokine cocktails at different concentrations on day 2 and day 5. (G) Bar plots illustrating percentages of CD3^-^ CD56^+^ and CD3^-^ CD33^+^ cells significantly increased with cocktail #2 at different concentrations on day 5, while percentages of CD3^+^ CD33^+^ and CD3^+^ CD56^+^ cells significantly increased with cocktail #2 at different concentrations on both day 2 and day 5.

To functionally validate the role of these identified microenvironmental factors in driving immune cell plasticity, we isolated and cultured CD3^+^ T cells from human primary PBMCs. Following treatment with individual cytokines or cocktails derived from the ligand-receptor analysis, cells were subjected to flow cytometry analysis on day 2 and 5 (Fig. 4B). The individual cytokines demonstrated limited efficacy in modulating cell plasticity, with only a subset showing any activity. For instance, IL15 and SAA1 induced CD3^+^ T cells to express more of the NK cell-specific marker CD56; TNFα induced CD3^+^ T cells to express the myeloid cell-specific marker CD14; and VEGFβ induced CD3^+^ T cells to express the progenitor-specific marker CD34 (Fig. 4C, D). The external factor cocktails were defined as follows: cocktail #1 (TGFβ1 and JAG1), cocktail #2 (IFNγ, IL15 and CXCL12), cocktail #3 (TNF and IGF1), and cocktail #4 (VEGFα and IL1B). Among these, we found that cocktail #2 exhibited a significant impact on T cell plasticity across all time points and concentrations (Fig. 4E and Fig. S8B, C). Flow cytometric analysis revealed that treatment with cocktail #2 induced distinct shifts in immune cell populations. The proportions of CD3^-^ CD56^+^ and CD3^-^ CD33^+^ cells on day 5 exhibited a statistically significant increase compared to the control group. Notably, the percentages of CD3^+^ CD56^+^ and CD3^+^ CD33^+^ cells were significantly elevated on both day 2 and day 5 (Fig. 4F, G). Collectively, our findings demonstrate that the cocktail induced a state of heightened plasticity in T cells, directing their transdifferentiation into NK-like or myeloid-like cells, thereby validating the role of the identified microenvironmental factors in driving the specific cell fate changes observed in our phylogenetic maps.

### Immune Cell Fate Change Accompanied by Epigenetic Reprogramming

To further validate the effect of cocktail #2, we treated human primary CD3^+^ T cells and performed multi-omics profiling (RNA-seq, ATAC-seq and WGBS (for DNA methylation)) across multiple time points (Fig. 5A). The RNA-seq revealed a clear dynamic transcriptional trajectory (Fig. 5B), while the ATAC-seq identified distinct patterns of chromatin accessibility between day 0 and day 8 (Fig. 5C). Furthermore, WGBS analysis demonstrated a global reduction in DNA methylation by day 8, evidenced by lower average beta values and an increase in hypomethylated loci (Fig. 5D, E). Collectively, our multi-omics profiling provided definitive evidence that cocktail #2 induced a profound reprogramming of T cell identity, thereby confirming its role in promoting cell plasticity.

**Fig. 5.**
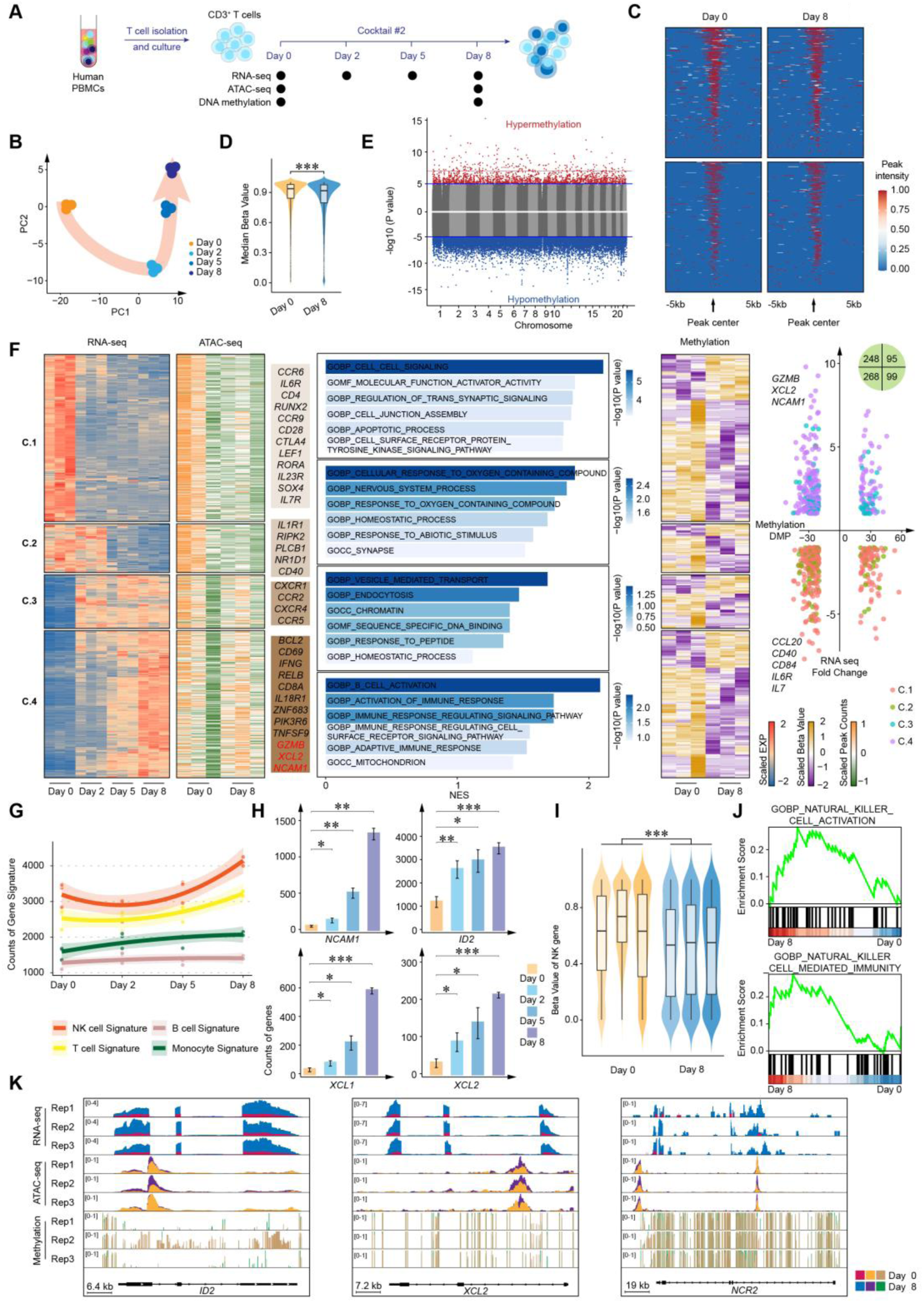
Cocktail induced DNA demethylation and upregulation of NK cell-associated genes in T cells. (A) Experimental diagram of human primary T cell stimulation with cocktail #2. RNA-seq was performed on day 0, day 2, day 5 and day 8. ATAC-seq was performed on day 0 and day 8. WGBS was performed on day 0 and day 8. (B) The PCA plot showing the temporal progression of the samples. (C) Tag heatmap of peak differences with cocktail #2 treatment on day 0 and day 8. (D) Violin plot showing the comparison of methylation levels in response to cocktail #2 on day 0 and day 8. (E) Scatter plot of hypomethylated and hypermethylated loci identified by comparative analysis of day 0 and day 8 methylation data. (F) Heatmap showing gene expression levels (RNA-seq), chromatin accessibility (ATAC-seq) and DNA methylation levels (WGBS) of the 711 overlapped genes. The scaled gene expression, scaled peak counts and scaled beta values were shown. Scaled expression of DEGs from RNA-seq analysis was used for K-means clustering analysis, with representative NK- or T cell-related genes labeled in each cluster. GSEA analysis were shown based each cluster. Scatter plot showing the distribution of the overlapped genes on RNA-seq fold change and methylation DMP. (G) Line graph depicting the dynamic changes of various RNA-seq signatures over time. (H) Bar plots showing the NK-related genes (*NCAM1*, *ID2*, *XCL1* and *XCL2*) significantly increased over time. (I) Violin plot showing the methylation levels of NK-related genes in response to cocktail #2 on day 0 and day 8. (J) GSEA analysis showing that demethylated promoters are associated with NK cell activation and differential peaks are associated with NK cell-mediated immunity, respectively. (K) Transcriptome (RNA-seq), chromatin accessibility (ATAC-seq) and DNA methylation (WGBS) profiles at the NK cell-related genes *ID2*, *XCL2* and *NCR2* locus in T cells after the cocktail #2 treatment.

Upon comparing sequencing data from day 0 and day 8, we identified differentially expressed genes (DEGs), differentially accessible regions (DARs), and differentially methylated positions (DMPs) (Fig. S9A). A total of 711 genes overlapped among these three sets, and GO enrichment analysis indicated their involvement in T cell differentiation and immune response (Fig. S9B, C). These overlapping genes were categorized into four clusters. We observed that the upregulation of DEGs in cluster 3 (C.3) and 4 (C.4) was accompanied by increased chromatin accessibility (Fig. 5F). In cluster 1 (C.1), several T cell-related genes—including *CCR6*, *IL6R* and *CD4*, which are involved in cell signaling—were consistently downregulated after cocktail treatment. Notably, in C.4, we detected upregulation of NK cell-associated genes such as *XCL2* and *NCAM1*. Additionally, genes encoding cytotoxic factors (*IFNG* and *GZMB*) were also activated, showing moderate upregulation in cocktail-induced NK-like cells compared to controls (Fig. 5F). Across all four clusters, hypomethylated genes significantly outnumbered hypermethylated ones, a finding consistent with previous reports of global DNA hypomethylation during cell reprogramming[30]. Based on this premise, we further classified promoters of DMPs between day 0 and day 8 into six clusters. Strikingly, in C.3, we found that NK cell-related genes including *KLRK1*, *NKG7*, *CD226* and *KLRC1* exhibited decreased DNA methylation and concurrent increases in gene expression (Fig. S9D). These NK cell-related genes were associated with NK cell-mediated immunity and regulation of NK cell-mediated immunity. Collectively, these results demonstrate that the cocktail treatment effectively reprogrammed T cells, leading to a transdifferentiation towards an NK-like phenotype, which was achieved through a coordinated series of changes at the transcriptional, epigenomic, and DNA methylation levels.

RNA sequencing revealed a time-dependent enhancement of the NK cell signature (Fig. 5G). Key NK cell-related genes, including *NCAM1*, *XCL1* and *XCL2*, along with the developmental transcription factor *ID2*, showed progressive upregulation over time (Fig. 5H). Cytotoxic factor genes *GZMA* and *GZMB* were also significantly induced. Additionally, natural cytotoxicity receptors *NCR1*, *NCR2* and *NCR3* gradually increased (Fig. S9E). While the overall T cell signature and *CD3D* were largely maintained, we observed a decline in *CD4* and a concurrent rise in *CD8A*, indicating that the emergence of NK-like cells was not attributable to a loss of T cell identity. This phenotypic shift was further supported by decreased DNA methylation at NK cell-related gene loci (Fig. 5I). GSEA linked promoter DMPs to NK cell activation and differential ATAC-seq peaks to NK cell immunity (Fig. 5J), and flow cytometry confirmed an increase in NK-like cells (Fig. S9F). We then analyzed integrated multi-omics profiling from T cells treated with cocktail to validate the coregulation of selected NK cell-related genes (Fig. 5K and Fig. S9H). The cocktail treatment activated the expression of *ID2*, *XCL2*, *NCR2*, *XCL1* and *NCR3*, which was concomitantly associated with increased chromatin accessibility and DNA hypomethylation at these gene loci. In contrast, *CD3D* expression remained largely stable across all three sequencing assays, whereas *CD4* was transcriptionally inactivated and exhibited reduced chromatin accessibility. (Fig. S9I). The T cell transcription factor *BCL11B* showed mild downregulation and hypermethylation, consistent with its reported role in restraining NK-like reprogramming[31]. Collectively, these data underscore that the cocktail #2 remodels the DNA methylome and transcriptome of T cells, driving their transition into NK-like cells.

### Immune Cell Dynamics Associated with Cancer Prognosis

To investigate whether there were specific occurrences of immune cell state transitions and fate changes within the tumor microenvironment, we examined these cells in normal bone marrow (BM)[32] and tissues under pathological conditions. In BM, there were hematopoietic stem cells and multiple progenitors, but few mature cells as previously reported (Fig. 6A). For these cells, most underwent expansion. Cell dedifferentiation occurred at low frequency, including erythrocyte into early-erythrocyte, and CD4^+^ T cell into naïve CD4^+^ T cell. Transdifferentiation mainly happened between CD4^+^ T cell and CD8^+^ T cell, and from CD4^+^ T cell to GMP and NK cell (Fig. 6B). In pathological ovarian tissues (labelled as NOS, nonmalignant samples), immune cell heterogeneity was comparable with that in cancer. A greater number of cell transdifferentiations were observed when compared to bone marrow data. State transition happened between CD8^+^ T GZMK^+^ Tex cell and CD8^+^ T Trm cell (Fig. 6B). When comparing BM, NOS, and high-grade serous ovarian cancer (HGSOC), immune cell expansion, differentiation, and transdifferentiation occurred at relatively higher frequencies compared to dedifferentiation and state transition. Furthermore, the frequency of transdifferentiation gradually increased from BM to NOS and HGSOC, with HGSOC exhibiting significantly higher rates than BM (Fig. 6C). Consequently, immune cell plasticity and dynamics are widespread *in vivo*, occurring at variable frequencies that are influenced by pathological and malignant conditions.

**Fig. 6.**
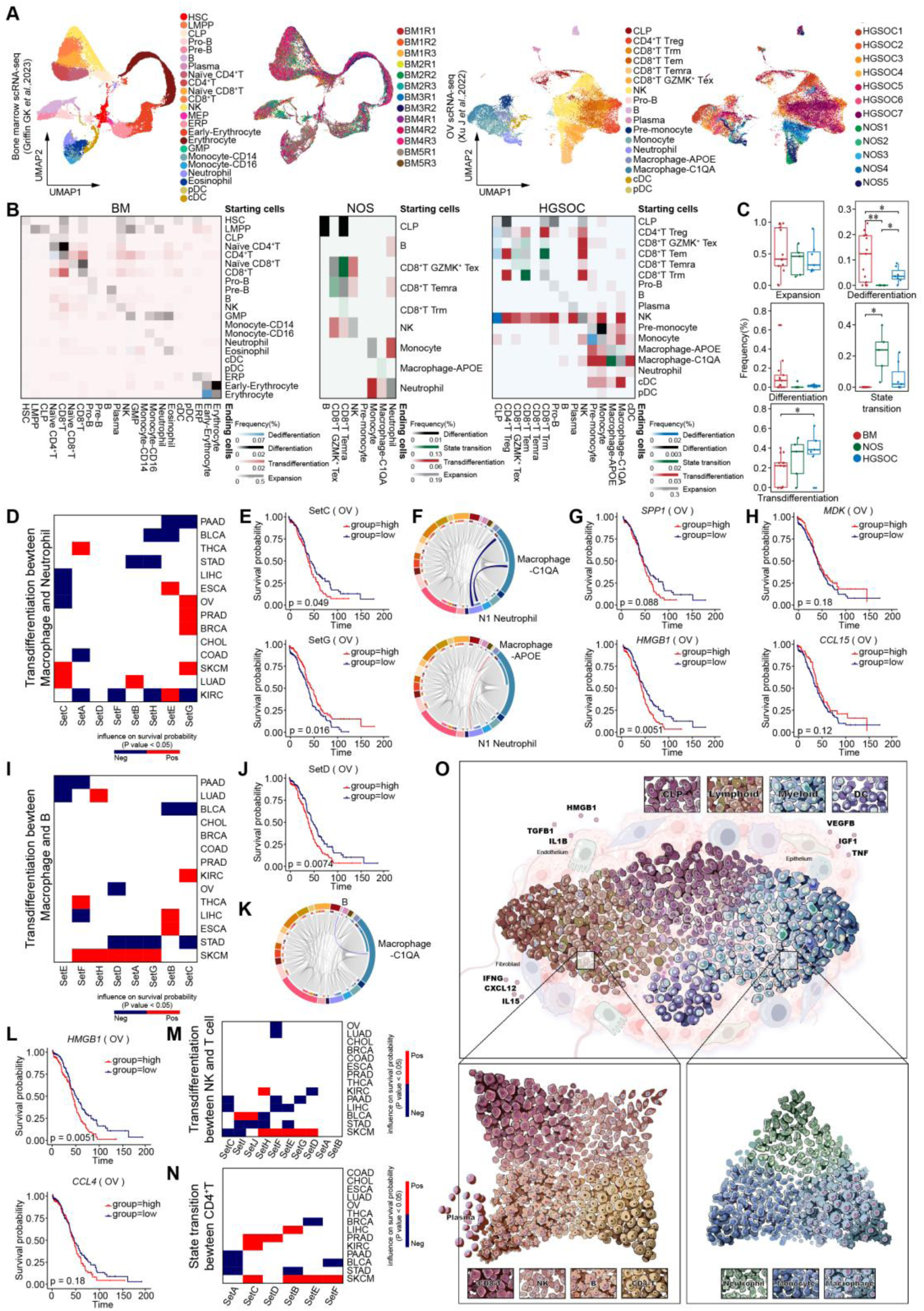
Correlation of immune cell dynamics to the clinical outcome. (A) UMAP plots showing the bone marrow scRNA-seq project (Griffen GK *et al*., 2023) and the ovarian scRNA-seq project (Xu J *et al*., 2023). UMAP plots displaying the distribution of each sample, respectively. (B) Heatmaps showing the frequency of cell change types in the BM (bone marrow), NOS (nonmalignant samples) and HGSOC (High-Grade Serous Ovarian Cancer), respectively. The cell types in the rows are starting cells, and the cell types in the columns are ending cells. Different colors represent different cell change types. The frequency is calculated through dividing the counts of each cell change type by the total number of all cell change types, respectively. (C) Box plots showing the proportions of the cell change types across BM, NOS and HGSOC projects. (D) Heatmap illustrating the influence of the different gene sets on survival probability across various cancer. The gene sets affecting the transdifferentiation between macrophage and neutrophil are listed in Fig. S8A and Table S3. (E) Kaplan-Meier plots showing worse clinical outcome in OV patients with the higher expression of gene set C and better clinical outcome in OV patients with the higher expression of gene set G using the project of tcga_pan_can_atlas_2018 in TCGA dataset. (F) Circus plot (up) listing the transdifferentiation between N1 neutrophil and macrophage-C1QA affected by gene set C existing in OV project in Fig. 2B. Circus plot (down) listing the transdifferentiation from macrophage-APOE to N1 neutrophil affected by gene set G existing in OV project in Fig. 2B. (G) Kaplan-Meier plots showing worse clinical outcome in OV patients with the higher expression of *SPP1* and *HMGB1*. The two genes probably are more effective in gene set C. (H) Kaplan-Meier plots showing better clinical outcome in OV patients with the higher expression of *MDK* and *CCL15*. The two genes probably are more effective in gene set G. (I) Heatmap illustrating the influence of the different gene sets on survival probability across the various tumor. The gene sets affecting the transdifferentiation between macrophage and B are listed in Fig. S8A and Table S3. (J) Kaplan-Meier plots showing worse clinical outcome in OV patients with the higher expression of gene set D using the project of tcga_pan_can_atlas_2018 in TCGA dataset. (K) Circus plot (left) listing the transdifferentiation from B to macrophage-C1QA affected by gene set D existing in OV project in Fig. 2B. (L) Kaplan-Meier plots showing worse clinical outcome in OV patients with the higher expression of *CCL4* and *HMGB1*. The two genes probably are more effective in gene set D. (M) Heatmap illustrating the influence of the different gene sets on survival probability across the various tumor. The gene sets affecting the transdifferentiation between NK and T are listed in Fig. 4A and Table S3. (N) Heatmap illustrating the influence of the different gene sets on survival probability across the various tumor. The gene sets affecting the state transition between CD4^+^ T are listed in Fig. S8B and Table S3. (O) Model for the cellular types of tumor immune cells and their genetic and tumor environmental determinants. The model shows cell changing among the major lineages (up), among the lymphoid cells (down left) and among the myeloid cells (down right). Intermediate cells are represented between diverse cell types and subtypes. Fibroblast, endothelium and epithelium are labelled as well as partial ligands highly probably affecting changing sent by them are listed.

Lastly, we aimed to investigate the potential relationship between these changes in immune cells and cancer prognosis. For classifying external factors that contribute to transdifferentiation between macrophages and neutrophils, we grouped these factors into different sets (Table S3). We found that different sets of genes in various cancer types were associated with distinct probabilities of survival (Fig. 6D). As an illustration, in the case of OV samples, the high expression of SetC genes that govern the transdifferentiation between N1 neutrophils and macrophage-C1QA significantly hindered survival rates, while the high expression of SetG genes responsible for the transdifferentiation from macrophage-APOE into N1 neutrophils significantly enhanced survival likelihood (Fig. 6E, F). Notably, representative genes for each set, such as *SPP1* and *HMGB1* in SetC, and *MDK* and *CCL15* in SetG, were illustrated (Fig. 6G, H). Furthermore, the expression of SetD genes, including *HMGB1* and *CCL4*, that contribute to B cell transdifferentiation into macrophage-C1QA showed a positive correlation with survival in OV patients (Fig. 6I-L). A parallel set of observations was made for SetF genes in OV influencing the transdifferentiation between NK cells and T cells (Fig. 6M). Additionally, all gene sets involved in state transitions among diverse CD4^+^ T subtypes in BRCA, PAAD, BLCA, and STAD patients exhibited a significant correlation with poor survival, while those in LIHC, PRAD, KIRC, and SKCM patients were associated with enhanced survival (Fig. 6N).

## Discussion

In this study, we generated phylogenetic trees for immune cells within multiple solid tumors based on SNP and mtDNA mutation inferred from scRNA-seq data, combined with cell type definitions. Our results demonstrate that immune cells could acquire plasticity and exist at highly dynamic states or stages. Generally, not only could cell subtypes within lymphoid or myeloid lineages transition to each other, but also the cell types could switch between each other. Comparing with other cells, B cells showed a relatively higher dynamic potential to change into multiple cell types, including T cells, macrophages and neutrophils, and can also be derived from these lineages. Conversely, DCs appear to maintain a more stable identity. Almost all these cell state transitions or fate changes require the initial acquisition of plasticity by becoming ICs with suppression of starting cell-specific genes and gradual establishment of an ending cell transcriptional signature. These changes were highly influenced by a mixture of tumor microenvironment extrinsic factors, such as CXCL12, IL15, TGFβ, TNF and IFNγ, which have been reported to affect cell fate, especially epithelial-mesenchymal-transition (Fig. 6O). We also found a cocktail can induce human primary T cells to acquire plasticity to potentially transdifferentiate into NK-like cells within the tumor microenvironment.

Previous studies have reported that immune cells within tumors or under pathological conditions could acquire plasticity, with a primary emphasis on subtype transition, particularly for neutrophils and macrophages with pro- and anti-inflammatory properties, classified as type I or type II[3–5]. T cells could change into NK-like cells by inhibition of BCL11B and DNMT1. Th17 could transdifferentiate into Treg cells by TGFβ modulating Smad3/AhR signaling during immune response[28]. Only a few reports have described cell changes between major clusters. In a breast cancer mouse model, CSF1R^+^Pax5^Low^ B-cell precursors transdifferentiated into macrophage-like cells, coaxed by cancer-secreted macrophage colony-stimulating factor[16]. In advanced cervical carcinoma and multiple animal tumor models, CD45^+^ erythroid precursor cells gradually acquired myelopoietic potential and finally transdifferentiate into erythroid differentiated myeloid cells under the influence of granulocyte-macrophage colony-stimulating factor[14]. Both induced cells lead to a decline in T cell-mediated tumor suppression. Beyond these sporadic cases, our study provides a systemic view of all possible immune cell phenotype changes in pan-cancer, not only verifying previous reports but also discovering previously unknown changes.

Cell fate changes have been widely, artificially achieved both *in vitro* and *in vivo* by modulating transcription factor expression for not just solid tissue cells, but also immune cells. Additionally, microRNAs and small molecules that regulate epigenetics and canonical stemness-related signaling pathways have been used successfully to induce cell changes[33–35]. In addition to these key driving forces, cell culture conditions, such as oxygen pressure[36], and contextual stiffness[37], as well as microenvironment soluble factors, such as growth factors and cytokines[38], all affect cell fate change direction and efficiency. Given all these inducing factors are present in solid tumors as a complex combination, it is unsurprising that so many immune cells acquire plasticity, undergo state transition and fate change, combined with widespread gene mutation in cancer. Investigating each cell fate change pair and the mechanism controlling the changing direction not only helps understand the immune cell heterogeneity and plasticity in cancer but also could enhance current immunotherapy[39].

The current gold standard for lineage tracing relies on genetic labelling through either fluorescent protein or genetic barcodes. Unfortunately, this method is limited to *in vitro* assays or animal models, making it challenging to utilize for human sample analysis[40]. Single-cell RNA and DNA sequencing analysis of the same cell is considered to be the optimal technique for lineage tracing analysis. However, this method is often expensive and low-throughput, which prevents widespread use[41, 42], Only genomic sequencing can provide a detailed genetic code that is necessary for constructing accurate phylogenetic trees, but losing cell type information. Multiple bioinformatic algorithms based on RNA sequencing, such as RNA velocity and pseudotime analysis, can help predict cell fate changes[43]. These methods, while valuable, offer predictive insights rather than definitively establishing ancestor-descendant relationships. The use of either SNP or mtDNA mutation alone, as previously explored, provides an incomplete picture of cellular lineage. Here, for the first time, we synergistically integrate both SNP and mtDNA mutation inferred from scRNA-seq data. This dual-genetic-code approach, combined with strict filtering criteria, substantially mitigates the noise and false-positive signals inherent to each individual method, thereby generating a far more precise and reliable phylogenetic tree. Beyond identifying previously reported immune cell type and subtypes changes, our approach revealed numerous novel inter-lineage relationships. Although these newly discovered cell changes require future confirmation through canonical lineage tracing methods, they provide a comprehensive atlas and framework that will guide subsequent mechanistic investigations.

## Acknowledgements

This work is supported by the National Key R&D Program of China (2024YFA1803503, 2023YFC2508900), the Shanghai Municipal Health Commission (2022XD050), and the Shanghai Municipal Education Commission (22SG12).

## MATERIALS AND METHODS

### Software and algorithms

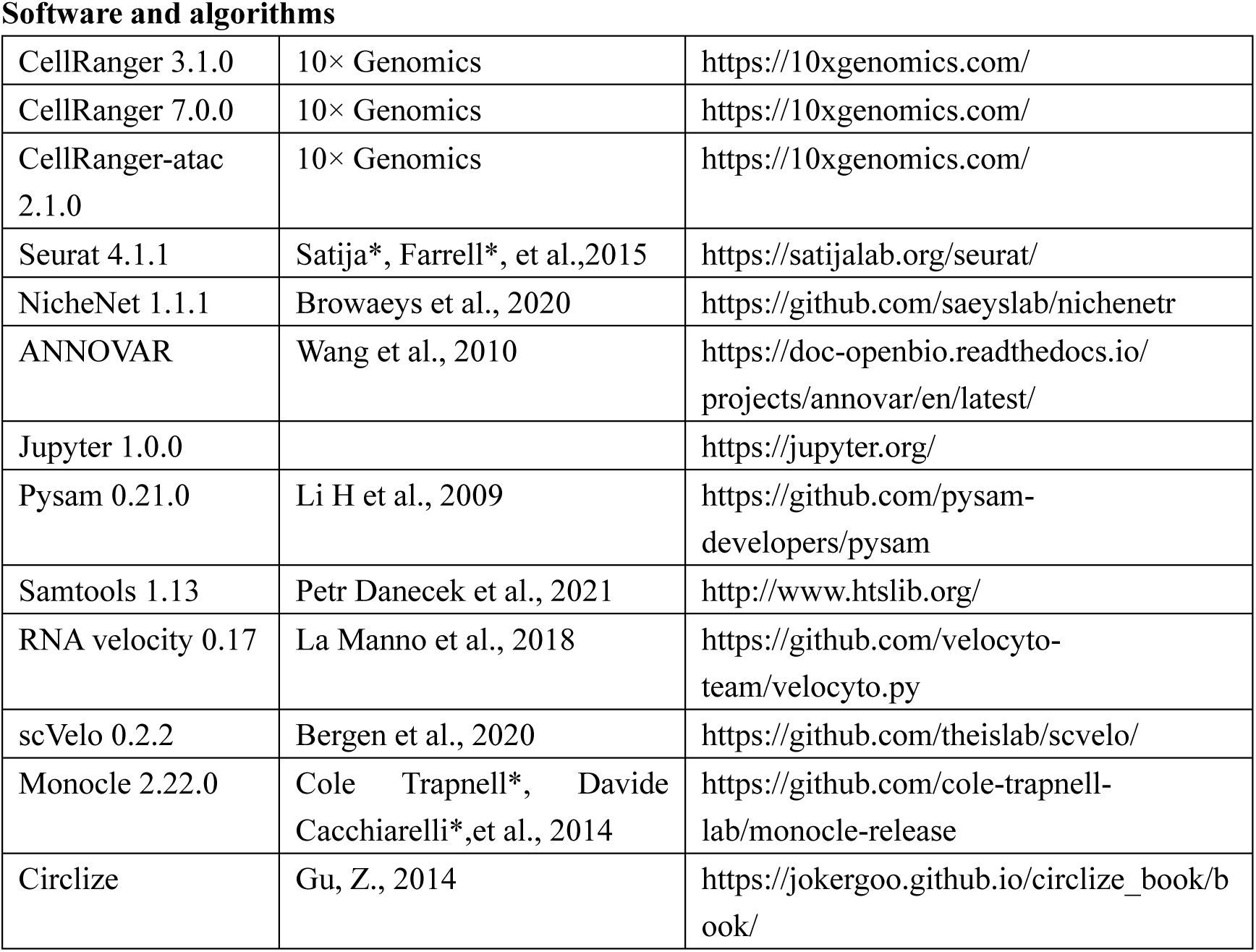
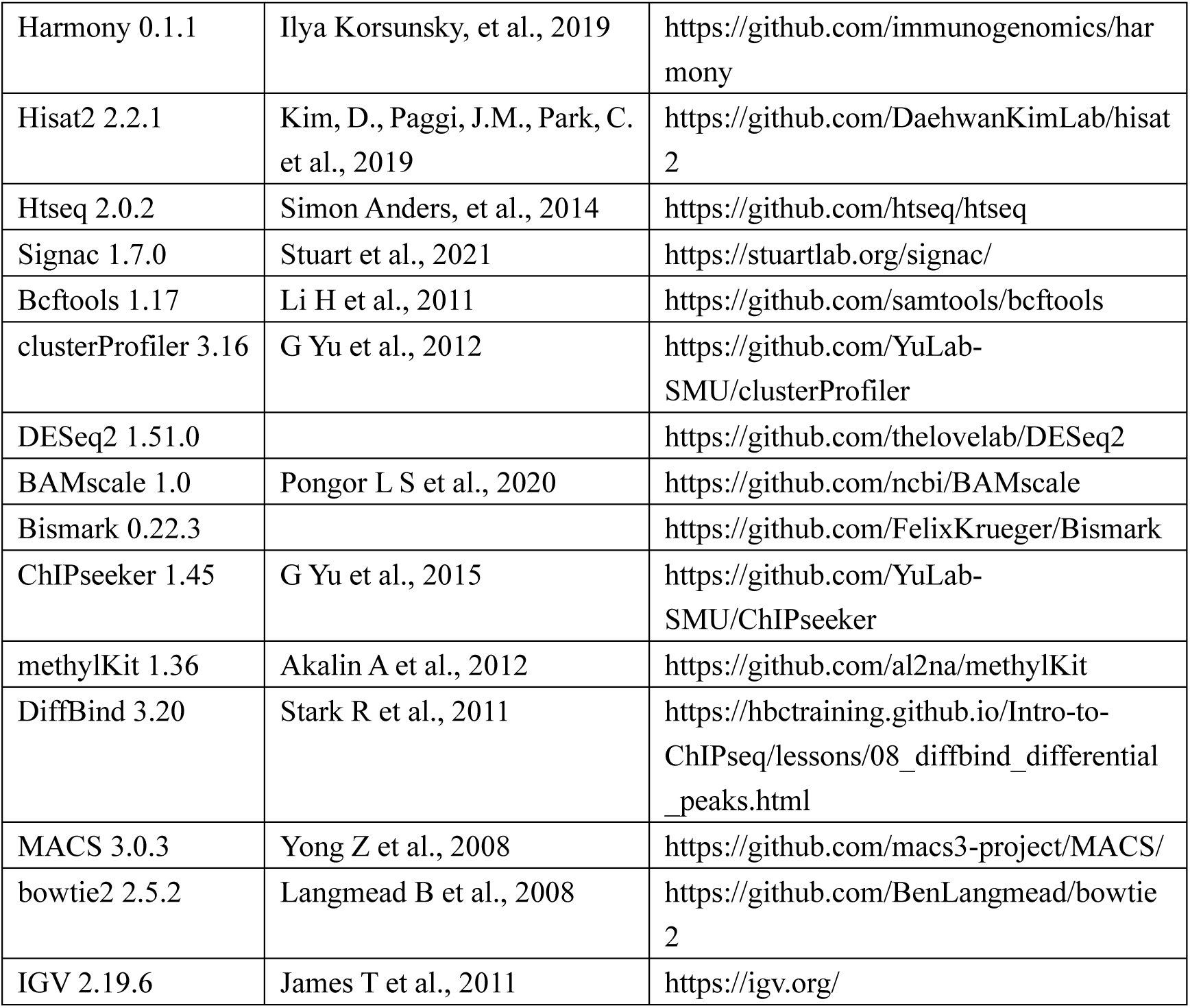

## METHOD DETAILS

### Single-cell RNA-Seq datasets collected in this study

We collected raw single-cell sequencing data on tumor-infiltrating hematopoietic cells from approximately 150 samples across 19 projects, which were diagnosed as one of the 14 common tumor types defined by the TCGA database (Fig. 2A and Table S1). Our team obtained the data by downloading them from the GEO, ENA, BioStudies, and GSA databases using SJTU HPC. The data included 10× scRNA-seq, 10× scATAC-seq, and SMART-seq. We only selected data that met the following criteria: it had sequenced information on both the tumor cells and infiltrated immune cells, and the raw sequence data or bam files were available for download. Bone marrow data, ovarian benign tissue data and TCR sequencing were also downloaded for the further verification and analysis.

## QUANTIFICATION AND STATISTICAL ANALYSIS

### Single-cell RNA-seq data processing

All upstream analysis processes for scRNA-seq data were carried out on the SJTU HPC platform. Fastq files or bam files were obtained from the Siyuan No.1 platform using the wget function, SRA toolkit, or Aspera software. Bam files were then converted to the fastq file format using the Cell Ranger (version 3.1.0) bamtofastq function. To obtain bam files and expression matrixs, we processed the fastq files through the 10× scRNA-seq Cell Ranger Count pipeline using the GRCh38 human reference genome. Only quality-passed cells that passed the Cell Ranger filtering step were included in downstream analysis.

### Single-cell ATAC-seq data processing

We obtained the 10× scATAC-seq data from the GEO database and analyzed them using the 10× Cell Ranger-atac pipeline (version 2.1.0) against the GRCh38 human reference genome to obtain bam files and the atac peak matrix. Only quality-passed cells that passed the Cell Ranger-atac filtering step were included in downstream analysis.

### Single-cell immune profiling

We obtained the 10× single cell immune profiling from the GEO206325. We analyzed the 5’ single cell gene expression using the 10× Cell Ranger pipeline (version 3.1.0) against the GRCh38 human reference genome to obtain bam files and the expression matrix, while we analyzed V(D)J transcripts and clonotypes for T cells using 10× Cell Ranger VDJ pipeline (version 7.0.0) against the GRCh38 human VDJ atlas reference. The output of the 5’ single cell gene expression is the same as that of the 3’ single cell gene expression. The output of the VDJ sequencing contains the contig annotations and the clonotypes. The TCR information was confirmed and filtered for the clonotype identification.

### SMART-seq data processing

The SMART-seq data were aligned using hisat2 software (version 2.2.1), which was indexed with the GRCh38 human reference genome. The output was in the form of sam files, which were then converted to sorted bam files using samtools (version 1.13). The expression matrixs were counted using htseq software (version 2.02). Only the quality-passed cells and matrixs were included in downstream analysis.

### Dimension Reduction and Unsupervised Clustering

The single-cell RNA expression matrixs underwent dimension reduction and unsupervised clustering using the Seurat pipeline (version 4.1.1) on RStudio at the HPC studio visualization platform. The cells were normalized and scaled for homogeneity, and highly variable genes were selected for downstream analysis. Principal component analysis (PCA) was performed on the scaled data to reveal the main axes of variation and denoise the data. Clustering of single cells by expression matrix was achieved by setting the parameter “resolution” of the FindClusters function to 2, generating more clusters for downstream cell type identification. Data were further reduced using the Uniform Manifold Approximation and Projection (UMAP) implemented in the RunUMAP function with default parameters. Cluster-specific marker genes were identified using the FindAllMarkers function with default parameters. To identify cell types, unsupervised clustering was performed, characterizing blood cells with the high-specific marker *PTPRC*. The second round of dimension reduction and unsupervised clustering was performed on these hematopoietic/immune cells to characterize the subsets of hematopoietic/immune cells, identifying lymphoid cells using *CD3D* and myeloid cells using *CD14* following the above strategy. Subdivided cell types were defined by more specific markers published in the study.

For single-cell ATAC peak matrixs, we used the Signac pipeline (version 1.7.0) on RStudio at the HPC studio visualization platform. First, we processed the chromatin data using the peak matrix and fragment file outputted by Cell Ranger-atac. We then performed data frequency-inverse document frequency (TF-IDF) normalization and selected the top peaks for dimensional reduction. Singular value decomposition (SVD) was performed on the TD-IDF matrix using the selected features (peaks). Data were further reduced using the Uniform Manifold Approximation and Projection (UMAP) implemented in the RunUMAP function with default parameters, as in the RNA analysis workflow. Gene activity matrixs were created for further cell type identification.

### Integration of Multiple scRNA-seq Datasets by Harmony

We performed two rounds of Harmony (version 0.1.1) analysis to integrate multiple scRNA-seq datasets and merge cells of the same type to eliminate batch effects caused by different projects within the 14 tumor types. In the first round, we merged samples from the same tumor type to remove batch effects resulting from diverse protocols and platforms, including the 10× Genomics platform and SMART-seq platform. The second round of Harmony analysis aimed to merge the 14 tumor types into a single dataset, further reducing batch effects. The output of the Harmony integration was used as the input data for downstream analysis.

### Detection of Mitochondrial Mutations by Pysam

The bam files generated by Cell Ranger were imported into the Pysam (version 0.21.0) module using a custom Python script. The “pysam.AlignmentFile” function was used to transfer the bam file into the alignment file, allowing us to conveniently iterate over all of the read mappings in a specified region. We paid particular attention to mitochondrial information, including the cell barcodes, position information of mitochondrial reads, sequenced base information, and the number of times the site was sequenced. We detected this information using the “pileup-engine” function, which allowed us to obtain the mitochondrial sequence data for the entire sample. To identify mitochondrial mutations, we aligned this sequence data with the human mitochondrial reference genome downloaded from the Ensemble website at the same mitochondrial position site. If the base we sequenced differed from the reference, we recorded it as a mutation. We then calculated the mitochondrial mutation frequency of the entire sample to facilitate lineage tracing and variant quality control. Allele frequency was calculated as the reads of a specific base at a position divided by the total reads at that position. The data of cells’ mitochondrial mutations were prepared for the next lineage tracing and further analysis.

### Detection of SNP by samtools and bcftools

To obtain SNP (Single Nucleotide Polymorphism) information for each cell, a series of steps were taken. First, the whole data bam was split into smaller bams for each cell using samtools. Then, samtools pileup was used to align the split bam files to the GRCh38 human reference genome. Next, bcftools (version 1.17) was employed to filter unreliable SNPs, which included those with a quality score of less than 10, a DP (read depth) score of less than 5, and SNPs near INDELs within 5 sites. The output files from this step were VCF (variant call format) files, which are a common file format for variant analysis. The VCF files for each cell were merged into a single VCF file for further analysis to trace the lineage of cells in the tumor. All of the above steps to obtain SNPs used the array job function on the HPC platform, which allowed scripts to run simultaneously.

### Prediction of cell relationships through mitochondrial mutations

To ensure accurate lineage tracing, we filtered out variants with an allele frequency less than 5% to eliminate those that may disrupt heteroplasmy due to sequencing errors or RNA editing. We had tested various cutoff values and found 5% to be appropriate for our analysis. Additionally, variants that occurred too infrequently cannot be used to predict cell relationships, which is our primary aim. After filtering, we obtained new mitochondrial mutation data for each sample, and we used a Python script with the ‘multiprocessing’ module to analyze the data in parallel and obtained results quickly. We computed the correlation between every two cells by building a matrix that contains their mitochondrial mutations and frequencies. To obtain preliminary cell change results, we filtered correlation values less than 0.8, which we found to be a convincing cutoff to produce reasonable and beneficial lineage tracing results. We hypothesize that downstream cells should have more mitochondrial mutations than upstream cells since decedent cells inherit the ancestor’s mutation points and accumulate additional mutations. Therefore, we calculated the mitochondrial mutation number of cell pairs and defined the cell with more variants as the downstream cell and the cell with fewer mutations as the upstream cell.

Additionally, we considered traditional cell change rules, which suggest that downstream cells contain mitochondrial mutations from upstream cells. Thus, we converted mitochondrial mutations into sets after filtering unreliable mutations. If one set was a subset of another set, we considered the corresponding cell of the first set as the upstream cell and the corresponding cell of the second set as the downstream cell. We used this cell relationship as a supplement to the results obtained in the previous step, which helped remove false positive cell pairs.

### Prediction of Cell Relationships by SNP

The VCF files from each single cell were merged into a single file to obtain a precise result of cell relationships. To reduce interference, we filtered the SNPs a second time based on their quality score, removing those with a score less than 30. Additionally, we filtered out SNPs with insufficient read counts, with cutoff values ranging from about 10 to 600 depending on the sample. We also considered the Variant Allele Fraction (VAF) to filter SNPs below a certain threshold.

Next, we ran the same python script as in the previous step to obtain the correlation values between every two cells. We obtained preliminary cell change results calculated by SNP after filtering out pairs with a correlation value less than 0.6. We arrived at this cutoff value after extensive testing, as it provided reasonable and beneficial results for lineage tracing.

Finally, we counted the number of SNPs in each cell to determine whether it is the upstream or downstream cell in the cell pair.

### Integration of the Two Relationships

To obtain a more accurate and precise result of cell change relationships, we merged the predictions obtained from mitochondrial mutation and SNP data. Since these two types of data provided complementary information, merging them increased the overall accuracy of the result. We also took into account the biological reality that a single cell can give rise to multiple offspring cells, but the reverse is not possible. Therefore, we excluded cases where multiple cells transferred into a single cell. However, we made an exception for the pair that had the highest correlation value calculated by either mitochondrial mutations or SNPs. After this merging process, we arrived at the ultimate relationships of hematopoietic/immune cells in the tumor. Across all samples and tumor types, we obtained a total of 9,137 pairs of hematopoietic/immune cells (Table S2).

### Matching both TCR clonotype and genetic variation lineage tracing

We used the output of 10× single cell immune profiling to get two relations. One of the relation was the TCR clonotype calculated by the TCR sequencing. The other relation was integrated by the mtDNA mutation lineage tracing and the SNP lineage tracing. We found some of the cell pairs matched both TCR clonotypes and genetic variation lineage tracing.

### Developmental Trajectory Inference by RNA Velocity and scVelo

To evaluate the developmental trajectory inference of hematopoietic/immune cells in the tumor, we utilized velocyto (version 0.17) to process the RCC samples’ output files from Cell Ranger, generating a loom file for further analysis. We then implemented the workflow of scVelo, which is the evolutionary version of RNA velocity, to calculate the direction of each cell. The resulting UMAP coordinates and arrow coordinates were combined to determine the cells’ change direction, which was classified into eight types for more accurate classification.

### Developmental Trajectory Inference by Monocle

Monocle2 (version 2.22.0) is a well-known tool used for pseudotime analysis to infer the developmental trajectory of cells. However, it is typically used to analyze physical cell development. In our study, we have applied the Monocle2 algorithm to characterize the developmental origins of hematopoietic cells in RCC.

### Lymphoid cell and Myeloid Cells Activity Program

To explore the cellular activity and tendencies in different tumor types, we compiled information on cell types into a single dataset, resulting in 286 groups of data with average expression values for the cells they contain. We then selected approximately 1600 genes, including over 900 lymphoid cell genes and over 500 myeloid cell genes, which are the top differential genes of the respective cell types, to calculate lymphoid cell gene scores and myeloid cell gene scores. Variance was then computed to represent the discreteness of each cell, which can provide insight into the cells’ tendencies to change. For lymphoid cells, the abscissa represents the lymphoid cell gene change tendency score, where a higher score indicates a greater likelihood of change, while the ordinate represents the lymphoid cell gene score, where a higher score suggests a greater resemblance to a lymphoid cell. For myeloid cells, the abscissa represents the myeloid cell gene score, where a higher score indicates a greater similarity to a myeloid cell, while the ordinate represents the myeloid cell gene change tendency score, where a higher score suggests a greater likelihood of change. For dendritic cells, which express both lymphoid cell genes and myeloid cell genes, the abscissa and ordinate represent their tendency towards lymphoid and myeloid cell gene change, respectively. Finally, for erythrocytes, which do not express lymphoid or myeloid cell genes and are not our focus, the abscissa represents the myeloid cell gene score, while the ordinate represents the lymphoid cell gene score. We repeated this process with different gene sets, including neutrophil gene set and macrophage gene set, as well as CD4^+^ T gene set and CD8^+^ T gene set, to explore changes in cells within the lymphatic and medullary systems, respectively.

### Intermediate co-expressor (IC) program

To investigate the intermediate cells of cell change, our goal is to identify these cells from the tumor hematopoietic/immune cells. Using the cell transdifferentiation from B cell to macrophage-APOE as an example, we defined a B cell gene set and a macrophage-APOE gene set. The intermediate cells must express both *CD19* and *APOE*, and we filtered the cells by requiring them to have both the B cell gene score and the macrophage-APOE gene score. As the B cells are the upstream cells, we set the B cell gene score as a negative number. The filtered cells’ abscissa is calculated as the sum of the B cell gene score and the macrophage-APOE gene score, while the ordinate is calculated as the B cell gene score divided by the macrophage-APOE gene score or the macrophage-APOE gene score divided by the B cell gene score, depending on which score is higher. As shown in Fig. 3B, cells that are located around the y-axis and have higher y scores are more likely to be intermediate cells. It’s important to note that different intermediate cells depend on different gene sets.

### NicheNet analysis

NicheNet (version 1.1.1) is a powerful tool for predicting the ligands that drive the transcriptomic changes of target cells. To identify potential ligands that influence cell change, we selected CD4^+^ T cells, CD8^+^ T cells, endothelial cells, epithelial cells, fibroblasts, and monocytes as sender cells, respectively, and used intermediate cells as well as new upstream cells as receptor cells. To evaluate the top active ligands, we chose the cells that showed expansion as the negative control cells and constructed interactions between ligands and receptors. We listed the top 40 ligands of each cell type and assigned scores to these top ligands corresponding to their ranks. The resulting dot plot shows the ligands involved in cell transdifferentiation between B cells and macrophages, the ligands of cell transdifferentiation between macrophages and neutrophils, the ligands involved in cell transdifferentiation between NK cells and T cells, and the ligands associated with the cell state transition of CD4^+^ T Th17 cells and CD4^+^ T Treg cells.

### GO analysis

Gene Ontology (GO) is a powerful tool for exploring the biological functions of genes, providing insight into the molecular function, cellular component, and biological process. To better understand the functional implications of the gene mutations we have identified, we employ ClusterProfiler (version 3.16) for analysis. This analysis is performed by comparing the gene output from annovar against the R packages ‘org.Hs.eg.db’.

### TCGA data analysis

The expression data of TCGA were downloaded. Overall survival (OS) from the TCGA Pan-Cancer Clinical Data Resource (TCGA-CDR) (Liu et al., 2018) were used to analyze patients’clinical outcomes. Specifically, to test the correlation of the cell change types and patients’ survival, we divided the genes in Fig. 6D, I, M and N into several gene set, respectively (Table S3). The mean expression of the gene sets ware used to group samples into high and low groups based on 14 tumor types respectively. For some specific genes, we used the real expression and divided samples into two groups.

### Multiplex Immunofluorescence

Multiplexed immunofluorescence staining (AiFang biological, # AFIHC024) was performed using a tyramide signal amplification (TSA)-based approach on formalin-fixed paraffin-embedded (FFPE) tissue sections, frozen sections, or cell smears. Briefly, after standard deparaffinization, rehydration, and antigen retrieval in pH 9.0 EDTA or pH 6.0 citrate buffer, endogenous peroxidase was blocked with 3% H₂O₂. Non-specific sites were blocked with normal goat serum or 3% BSA. Sections were sequentially incubated with a primary antibody, a corresponding poly-HRP-conjugated secondary antibody, and a TYR-520/570/690 TSA fluorophore, followed by antibody elution using either heat-mediated antigen retrieval buffer or a dedicated antibody elution solution. This cycle of staining and elution was repeated to label multiple targets with distinct TSA fluorophores. Finally, nuclei were counterstained with DAPI, and slides were mounted with anti-fade medium for imaging via fluorescence microscopy, confocal microscopy, or a multispectral imaging system. Paraffin-embedded tissue sections for immunofluorescence staining were obtained from a cohort including LIHC, SKCM, BLCA, OV, COAD, BRCA, PAAD, STAD, LUAD, RCC and PRAD samples.

### Isolation and Expansion of Human CD3^+^ T Cells

Purified human periphery blood mononuclear cells (PBMCs) from healthy donors were sourced from Ruijin Hospital. Briefly, PBMCs were isolated through density gradient centrifugation using Ficoll-Paque PLUS (Cytiva, #17144003). Blood samples were diluted at 1:1 ratio with DPBS then layered above 15 mL Ficoll-Paque PLUS in a 50 mL conical tube. Once centrifuged for 30 min at 400×g, the white layers of PBMCs were collected and washed. Using EasySep Human CD3 Positive Isolation kit (StemCell Technologies, #17751), the CD3^+^ cell fraction was collected following the manufacturer’s guidelines. CD3^+^ T cells were cultured in X-VIVO media, consisting of X-VIVO 15 medium (Lonza, Cat #04-418Q) with 10% heat-inactivated fetal bovine serum (BIOAGRIO, #S1370), 1× penicillin–streptomycin (Thermo Fisher Scientific, #15140122), and 200U/mL human IL-2 (PeproTech, #200-02) at 1×106 cells/ml. Cells were stimulated with Dynabeads Human T-Activator CD3/CD28 (Gibco, #111.31D) at a bead-to-cell ratio of 1:4 and incubate in a 5% CO2 incubator at 37°C for 7-10 days. When the cell density exceeds 2.5×106 cells/ml, split cultures back to a density of 1×106 cells/ml in fresh media. Remove the Dynabeads prior to following applications.

### Cytokine induction of Human CD3^+^ T Cells

Count the cells and resuspend to a density of 1×106 cells/ml in X-VIVO media supplemented with cytokines, and incubate in a 5% CO2 incubator at 37°C for 8 days. In the single cytokine induction experiment, the media was supplemented with 10/50/100 ng/mL APOE (UA BIOSCIENCE, #UA030062), APP (Sino Biological, #10703-H02H), CCL2 (Primegene, #204-02), CCL3 (Primegene, #204-03), CCL4 (Primegene, #204-04), M-CSF (UA BIOSCIENCE, #UA040016), CXCL12α (Primegene, #201-12a), CXCL14 (Absin, #abs04139), CXCL16 (Primegene, #201-16), FGF2 (UA BIOSCIENCE, #UA040007), HMGB1 (R&D Systems, #1690-HMB), ICAM-1 (UA BIOSCIENCE, #UA010179), IFN-γ (UA BIOSCIENCE, #UA040066), IFG-1 (UA BIOSCIENCE, #UA040055), IL-15 (UA BIOSCIENCE, #UA040010), IL-1β (Primegene, #101-01B), IL-6 (UA BIOSCIENCE, #UA040052), IL-33 (UA BIOSCIENCE, #UA040036), JAG1 (Absin, #abs04386), LGALS3 (Primegene, #603-03), MIF (UA BIOSCIENCE, #UA040119), NECTIN1 (UA BIOSCIENCE, #UA010017), SAA1 (Primegene, #602-26), SPP1 (UA BIOSCIENCE, #UA010434), TGF-β1 (UA BIOSCIENCE, #UA040085), TGF-β3 (Acrobio, #TG3-H5213), TNF-α (UA BIOSCIENCE, #UA040062), VEGF-121(UA BIOSCIENCE, #UA040082) or VEGF B (Sino Biological, #10544-H01H). In the cytokine combinations induction experiment, the media was supplemented with 100/200/300 ng/mL cytokines. All cytokines were human recombinant.

### Flow Cytometry

1×106 cultured cells were collected and washed once in 1×DPBS. Resuspend cells at 1 ml 1×DPBS and add 1μl Fixable Viability Stain 510 Stock Solution (BD Pharmingen, #564406), vortex immediately and incubate the mixture at 2-8°C for 30 minutes. Wash cells with Stain Buffer (FBS) (BD Pharmingen, #554656) and resuspend the cells in 50μl Brilliant Stain Buffer (BD Pharmingen, #563794). For cell surface markers, cells were incubated with antibodies at 4℃ for 30 min. For intracellular staining, cells were fixed and permeabilized with Fixation/Permeabilization kit (BD Pharmingen, #554714), then incubated with antibodies for 30 min. The following antibodies were used: APC-H7 anti-human CD3 (Clone SK7, BD Pharmingen, #560176), BUV395 anti-human CD11b (Clone ICRF44, BD Pharmingen, #563839), BV786 anti-human CD14 (Clone M5E2, BD Pharmingen, #563698), BUV737 anti-human CD16 (Clone 3G8, BD Pharmingen, #612786), BV650 anti-human CD19 (Clone SJ25C1, BD Pharmingen, #563226), PE anti-human CD33 (Clone WM53, BioLegend, #343404), FITC anti-human CD34 (Clone 561, BioLegend, #343604) R718 anti-human CD56 (Clone B159, BD Pharmingen, #566965), BV421 anti-human CD68 (Clone Y1/82A, BD Pharmingen, #564943), PE-Cy7 anti-human CD235a (Clone HI264 BioLegend, #349112). Samples were detected by BD FACSymphony A3. FlowJo v 10.8.1 was used for data analysis.

### Bulk RNA sequencing

Total RNA was extracted with Trizol Reagent (Invitrogen, #15596026CN), and its concentration, quality, and integrity were assessed using a NanoDrop spectrophotometer (Thermo Scientific). Starting from 3 μg of total RNA, mRNA was enriched using poly-T oligo-attached magnetic beads. The purified mRNA was fragmented in Illumina proprietary buffer with divalent cations at elevated temperature. First-strand cDNA was synthesized using random hexamer primers and SuperScript II reverse transcriptase, followed by second-strand cDNA synthesis with DNA Polymerase I and RNase H. The resulting cDNA fragments were end-repaired, adenylated at the 3′ ends, and ligated to Illumina PE adapters. Library fragments of 400–500 bp were size-selected using the AMPure XP system (Beckman Coulter, Beverly, CA, USA) and amplified with Illumina PCR Primer Cocktail in 15 cycles of PCR. The final library was quantified with the Agilent high sensitivity DNA assay on a Bioanalyzer 2100 system (Agilent Technologies) and sequenced on an Illumina NovaSeq 6000 platform at Shanghai Personal Biotechnology Co., Ltd.

### Whole-genome methylation sequencing

Genomic DNA (gDNA) was extracted using a commercial DNA extraction kit (Takara, #9765) according to the manufacturer’s protocol. DNA concentration and purity were assessed using a NanoDrop ND-2000 spectrophotometer (Thermo Scientific, USA). For library preparation, 100–200 ng of gDNA was spiked with 1% λDNA as an internal control and fragmented by sonication. The sheared DNA was size-selected, followed by bisulfite conversion to deaminate unmethylated cytosines into uracils. The bisulfite-treated DNA was then denatured into single strands, and an “A” base was added to the 3’ ends for adapter ligation. Adapter-ligated fragments were enriched by PCR to construct the final sequencing library. Library quality was evaluated using the Agilent High Sensitivity DNA assay on a Bioanalyzer 2100 system (Agilent Technologies). Sequencing was performed on an Illumina NovaSeq 6000 platform at Shanghai Personal Biotechnology Co., Ltd.

### ATAC sequencing

ATAC-seq libraries were prepared using the ATAC-Seq Kit (Active Motif, #53156) according to the manufacturer’s protocol. Briefly, 1×10⁶ cultured cells were collected, washed once with 1× DPBS, and lysed in 200 μL Lysis Buffer for 10 min on ice. After centrifugation, nuclei were washed with 1 mL Washing Buffer and collected by centrifugation at 500 ×g for 10 min at 4°C. Approximately 50,000 nuclei were used as input for library construction with the Hyperactive ATAC-Seq Library Prep Kit for Illumina. The nuclei were resuspended in 50 μL transposition mix and incubated at 37°C for 30 min, followed by immediate cooling at 4°C. Transposed DNA was purified using 2× DNA clean beads and amplified with P5 and P7 primers for 15 PCR cycles. Library quality was assessed by concentration measurement and fragment size distribution analysis. Final libraries were sequenced on an Illumina NovaSeq X Plus platform at Shanghai Personal Biotechnology Co., Ltd.

### RNA-seq analysis

The quality-controlled clean reads were then aligned to the human reference genome (hg38) using hisat2 (v2.2.1). The resulting BAM files were sorted and indexed using SAMtools (v1.13). Gene-level counts were generated by quantifying the number of reads mapped to each gene feature in the GENCODE annotation (v41) using htseq (v2.0.2). Differential expression analysis was performed on the raw count matrix using the DESeq2 (v1.51.0) package in R, with genes exhibiting an adjusted p-value (FDR) < 0.05 and |log2(FoldChange)| > 1 considered statistically significant.

### ATAC-seq analysis

The clean reads were then aligned to the reference human genome (GRCh38/hg38) using Bowtie2 (v2.5.2) with parameters tuned for ATAC-seq data (--very-sensitive -X 2000). Duplicate reads, mitochondrial reads and reads mapping to blacklisted regions were removed. Properly paired, non-duplicate reads were retained for downstream analysis. Peak calling was performed using MACS2 (v3.0.3). Differential accessibility analysis was conducted by comparing read counts in consensus peak regions across conditions using the DiffBind (v3.20) R package. Annotation and genomic distribution of peak regions relative to gene features were performed using the ChIPseeker (v1.45) R package.

### DNA methylome analysis

The clean reads were then aligned to the bisulfite-converted reference genome (GRCh38) using Bismark (v0.22.3), and duplicate reads were removed using the deduplicate_bismark script. Methylation calls were extracted using the bismark_methylation_extractor function. Differential methylation analysis was performed on the methylated and total read counts using the methylKit (v1.36) R package. Genomic regions with a methylation difference (delta) > 0.20 and an FDR < 0.05 were defined as significantly differentially methylated regions (DMRs). Annotation and genomic distribution of DMRs relative to gene features were performed using the ChIPseeker (v1.45) R package.

### Visualization of Genomic Data

Genomic tracks, including sequencing coverage (BAM files) and genomic intervals (BED or BigWig files), were visually inspected using the Integrative Genomics Viewer (IGV) (version 2.19.6). Specific loci of interest, such as differentially expressed genes from RNA-seq, differentially accessible regions from ATAC-seq or differentially methylated regions from WGBS were navigated to for detailed examination.

---

**Fig. S1.**
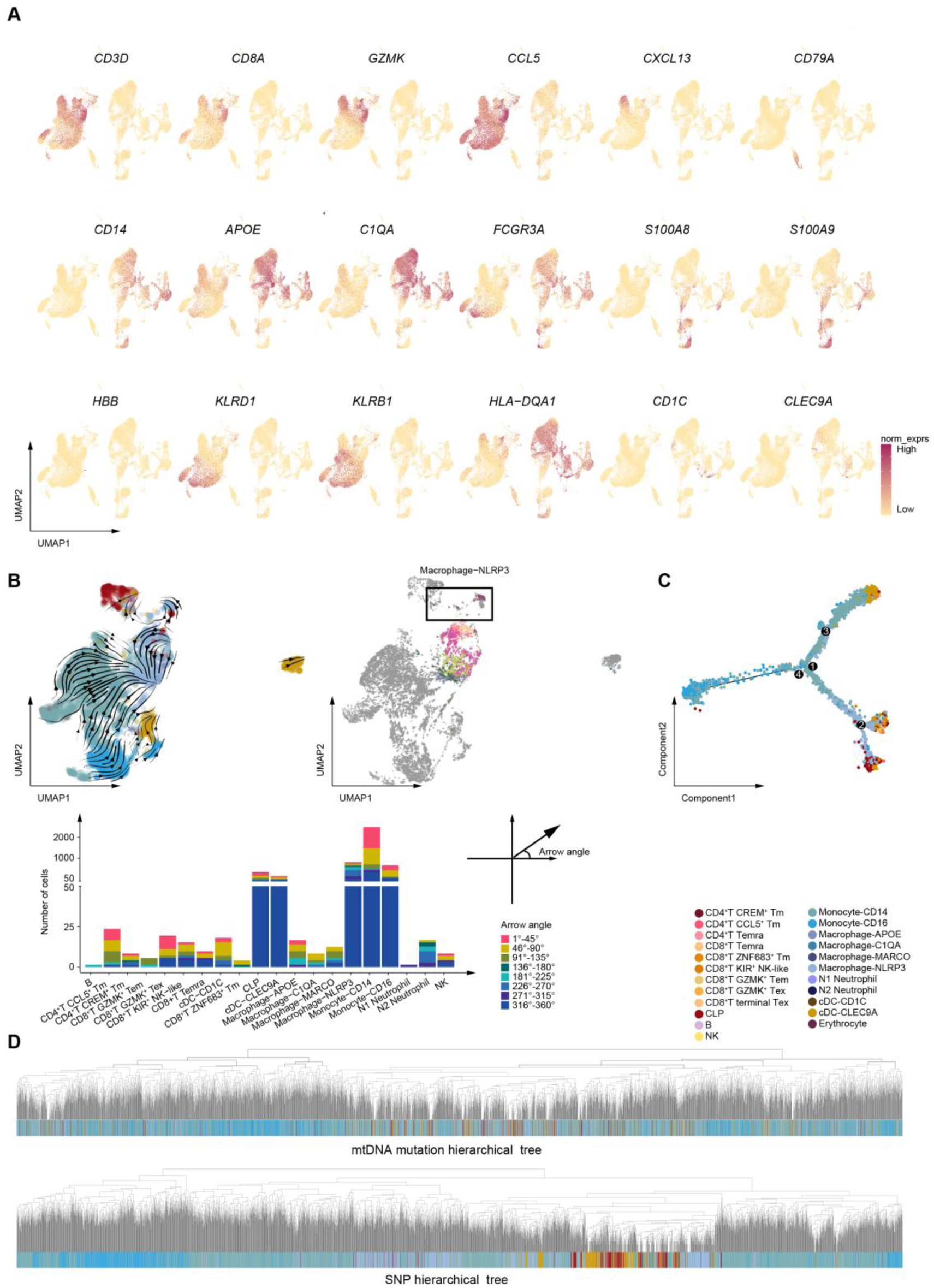
Gene expression and traditional lineage trajectory methods. (A) UMAP plots displaying the marker gene expression for the major lineages of immune cells, including T cells (*CD3D*, *CD8A*, *GZMK*, *CCL5*, *CXCL13*), B cells (*CD79A*), monocytes (*CD14*, *FCGR3A*), macrophages (*APOE*, *C1QA*), neutrophils (*S100A8*, *S100A9*), erythrocytes (*HBB*), NK cells (*KLRD1*, *KLRB1*), and DCs (*HLA-DQA1*, *CD1C*, *CLEC9A*). (B) RNA velocity analysis of the P1 pRCC sample (upper left). The cell type of macrophage-NLRP3 is highlighted by different colors according to the direction of the arrows (upper right). The black box displays the intricate directions of the macrophage-NLRP3 cells. Segmented bar chart shows the number of different arrow directions of the cell type. The arrow directions are divided into 8 types according to the arrow angle. (C) Developmental trajectory of immune cells inferred by Monocle2 as in (B). (D) A hierarchical tree of the mtDNA mutations (top) and the chromosome SNPs (down).

**Fig. S2.**
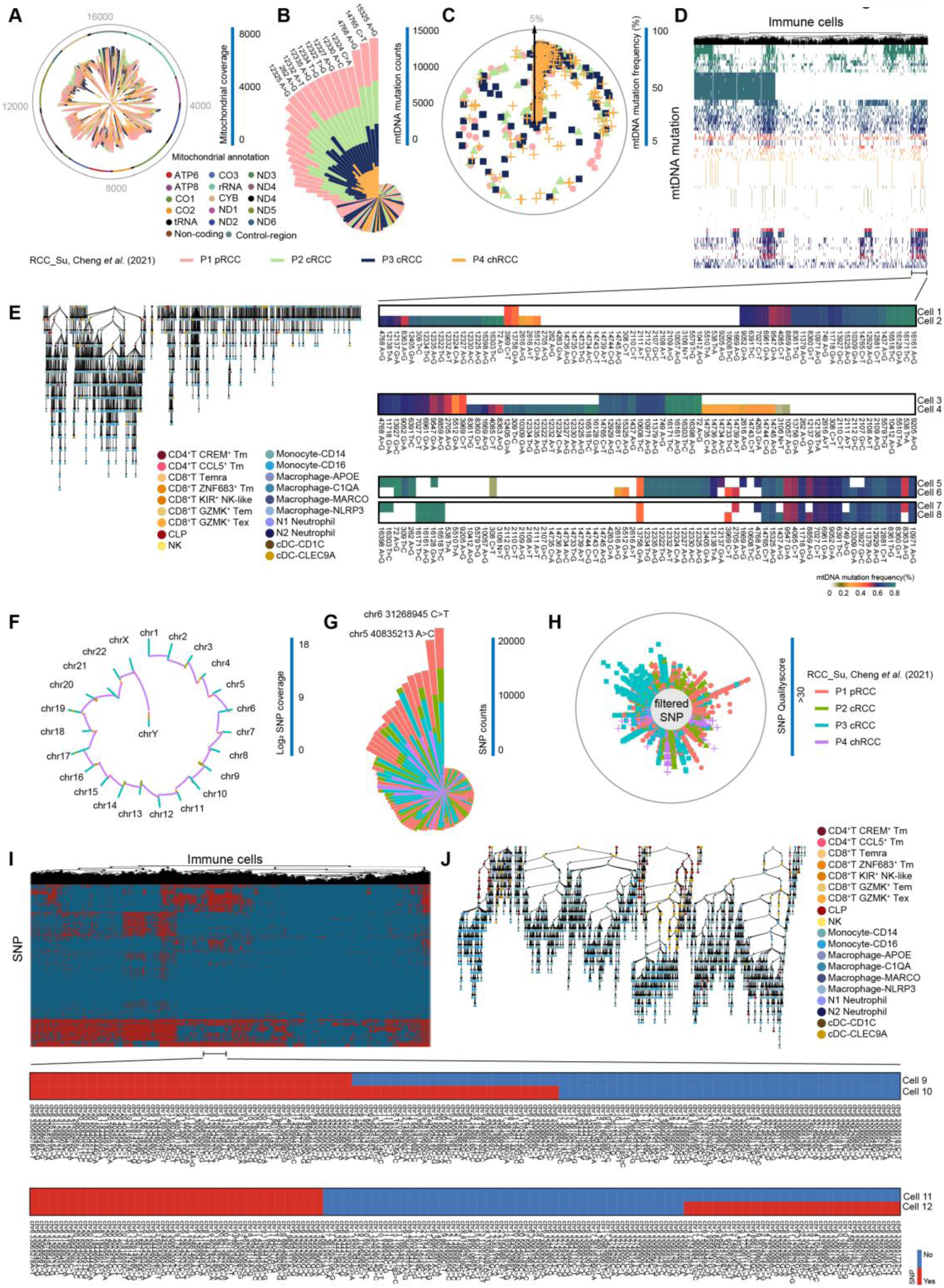
Lineage tracing through mtDNA mutation and SNP on different single-cell sequencing. (A) Coverage of the mitochondrial genome of the samples (RCC_Su, Cheng et al., 2021). Inner circle, coverage; middle circle, mitochondrial genome; outer gray circle, genome coordinates. Coverage is the sum of single cells. Panels (A) and (B) share the same legend as panel (C). (B) Nightingale Rose plots displaying the counts of mtDNA mutations across the four samples, with top mutations labelled. Panels (A) and (B) share the same legend as panel (C). (C) The frequency of mtDNA mutations across the four samples. Mutations whose frequency is less than 5% are filtered for the following lineage tracing. (D) Heatmap displaying the frequency of mtDNA mutations and hierarchical tree of immune cells in the P1 pRCC sample (RCC_Su, Cheng et al., 2021). The phylogram (up) represents cell-cell relationships, and the matrix (down) represents the different types of cell-cell relationships. The cell 1-cell 2 relationship and the cell 3-cell 4 relationship represent the mtDNA mutations of the descendant cells, which are cell 2 and cell 4, do strictly include those of the ancestor cells, which are cell 1 and cell 3. The cell 5-cell 6 relationship and the cell 7-cell 8 relationship represent the mtDNA mutations of the descendant cells, which are cell 6 and cell 8, do not strictly include those of the ancestor cells, which are cell 5 and cell 7. (E) A up-down phylogenetic tree calculated by mtDNA mutations showing the cell change process, meaning the ancestor cells transfer to the descendant cells (from up to down) in the P1 pRCC sample (RCC_Su, Cheng et al., 2021). Every dot represents a cell whose color indicates the cell type. (F) Coverage of the chromosome SNPs of the samples (RCC_Su, Cheng et al., 2021). Coverage is the sum of single cells. Panels (F) and (G) share the same legend as panel (H). (G) Nightingale Rose plots displaying the counts of chromosome SNPs across the four samples, with top SNPs labelled. Panels (F) and (G) share the same legend as panel (H). (H) The quality score of chromosome SNPs across the four samples. SNPs whose quality score is less than 30 are filtered for the following lineage tracing. (I) Heatmap displaying the chromosome SNPs and hierarchical tree of immune cells in the P1 pRCC sample (RCC_Su, Cheng et al., 2021). The phylogram (up) represents cell-cell relationships, and the matrix (down) represents the detailed information of cell-cell relationships. The cell 9-cell 10 and cell 11-cell 12 relationships represent the chromosome SNPs of the descendant cells (cell 10 and cell 12) do strictly include those of the ancestor cells (cell 9 and cell 11). The relationships that the chromosome SNPs of the descendant cells do not strictly include those of the ancestor cells are not displayed. (J) A up-down phylogenetic tree calculated by the chromosome SNPs showing the cell change process, meaning the ancestor cells transfer to the descendant cells (from top to bottom) in the P1 pRCC sample (RCC_Su, Cheng et al., 2021). Every dot represents a cell whose color indicates the cell type.

**Fig. S3.**
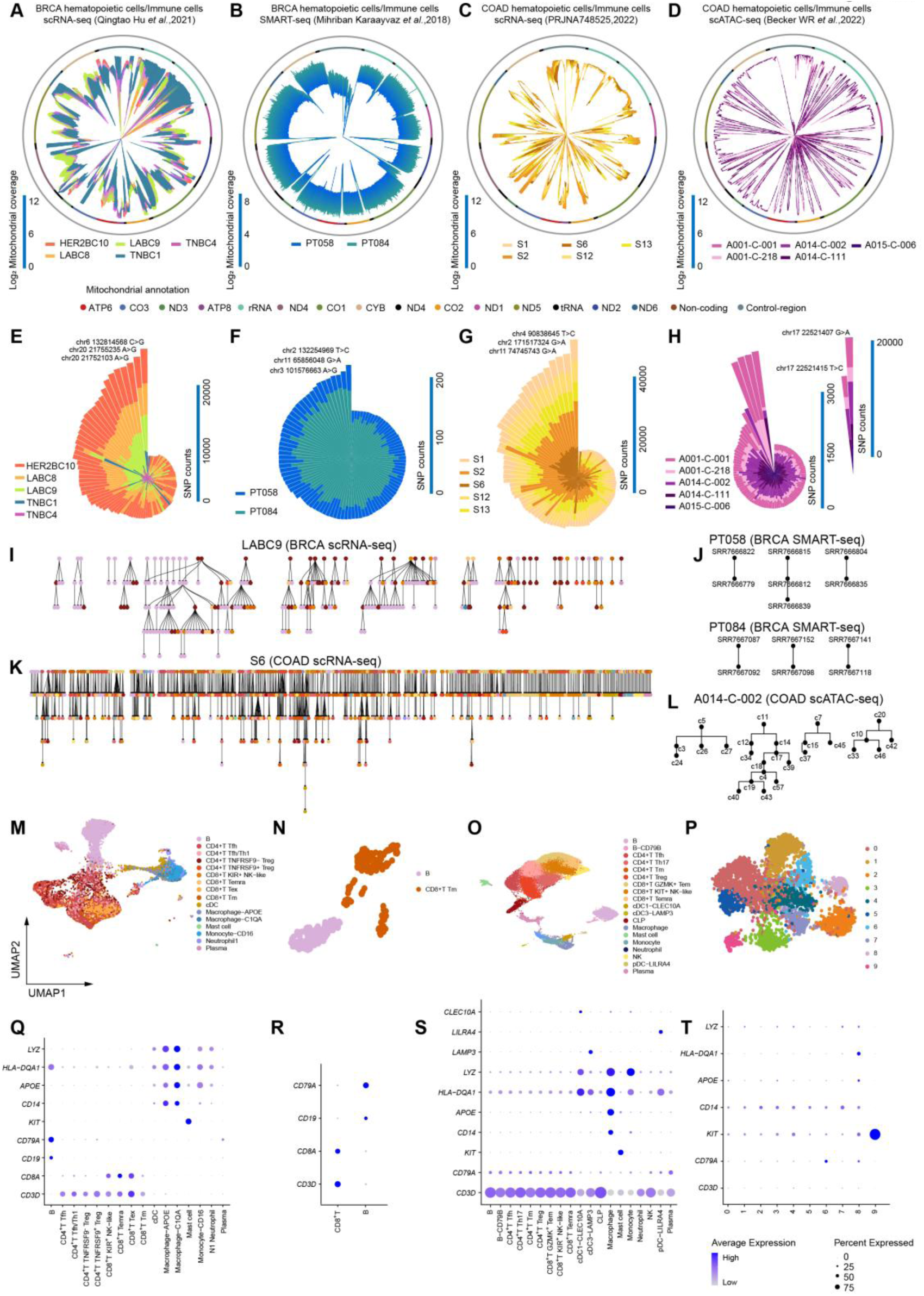
Comparison of lineage tracing methods in this study on different single-cell sequencing. (A, B, C, D) Coverage of the mitochondrial genome in the 10x scRNA-seq BRCA samples (Qingtao Hu et al.,2021); the SMART-seq BRCA samples (Mihriban Karaayvaz et al., 2018); the 10x scRNA-seq COAD samples (PRJNA748525, 2022) and the 10x scATAC-seq COAD samples (Becker WR et al., 2022),respectively. Inner circle, log2 coverage; middle circle, mitochondrial genome; outer gray circle, genome coordinates. Coverage is the sum of single cells. (E, F, G, H) Nightingale Rose plots showing the counts of chromosome SNPs across the samples in (A), (B), (C) and (D), respectively. The top SNPs are labeled. (I, J, K, L) Phylogenetic trees, built by both mtDNA mutations and SNPs, showing the cell change process, meaning the ancestor cells transfer to the descendant cells (from top to bottom) across the LABC sample in (A), the PT058 and PT084 sample in (B), the S6 sample in (C) and the A014-C-002 sample in (D), respectively. Every dot represents a cell whose color indicates the cell type. The colors of cell type correspond to the cell type in Fig. S5. (M, N, O, P) UMAP plots showing the four sequencing data as in (A), (B), (C) and (D), respectively. (Q, R, S, T) Bubble plots showing the marker gene expression of the major cell types across the samples in (A), (B), (C) and (D), respectively.

**Fig. S4.**
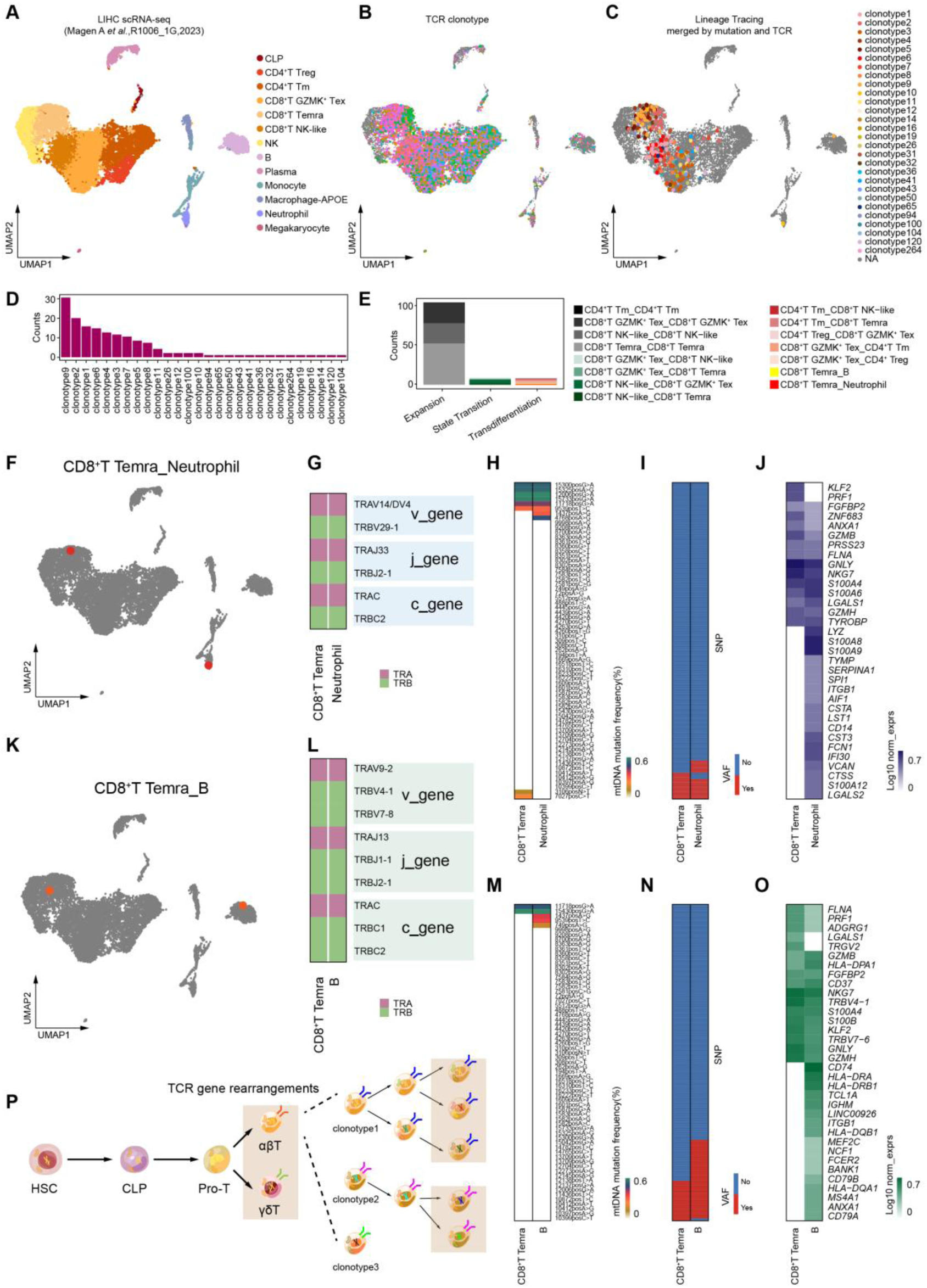
Identification diverse phenotypic states between TCR repertoire and genetic variation. (A) UMAP plots showing the hematopoietic/immune cells across the R1006_1G sample in the 10× scRNA-seq LIHC project (Magen A et al.,2021). (B) UMAP plots showing the clonotypes identified with paired TCR sequencing. (C) UMAP plots showing the cell pair identified with both paired TCR sequencing and genetic variation lineage tracing method. (D) Bar plot illustrating the counts of the clonotypes. (E) Bar plot illustrating the counts of the three cell change types. (F) UMAP plots showing the cell pair of CD8^+^ T Temra and Neutrophil. The cell pair is identified with both paired TCR and genetic variation. (G) Heatmap reporting the TCR gene in α chain and β chain of the cell pair in (F). (H) Heatmap showing the mtDNA mutation of the cell pair in (F). (I) Heatmap showing the SNP of the cell pair in (F). (J) Heatmap showing the transcriptional expression of the cell pair in (F). (K) UMAP plots showing the cell pair of CD8^+^ T Temra and B cell. The cell pair is identified with both paired TCR and genetic variation. (L) Heatmap reporting the TCR gene in α chain and β chain of the cell pair in (K). (M) Heatmap showing the mtDNA mutation of the cell pair in (K). (N) Heatmap showing the SNP of the cell pair in (K). (O) Heatmap showing the transcriptional expression of the cell pair in (K). (P) Pattern diagram showcasing the progress of T cells’ differentiation and the variation trend of TCR and genetic variation. Same mitochondria and chromatin mean that cells have similar genetic variation and different TCR till TCR gene arrangements. After TCR gene arrangement, T cells become different clonotypes so that different colors of mitochondria and chromatin and same color of TCR mean T cells have same TCR and different genetic variation.

**Fig. S5.**
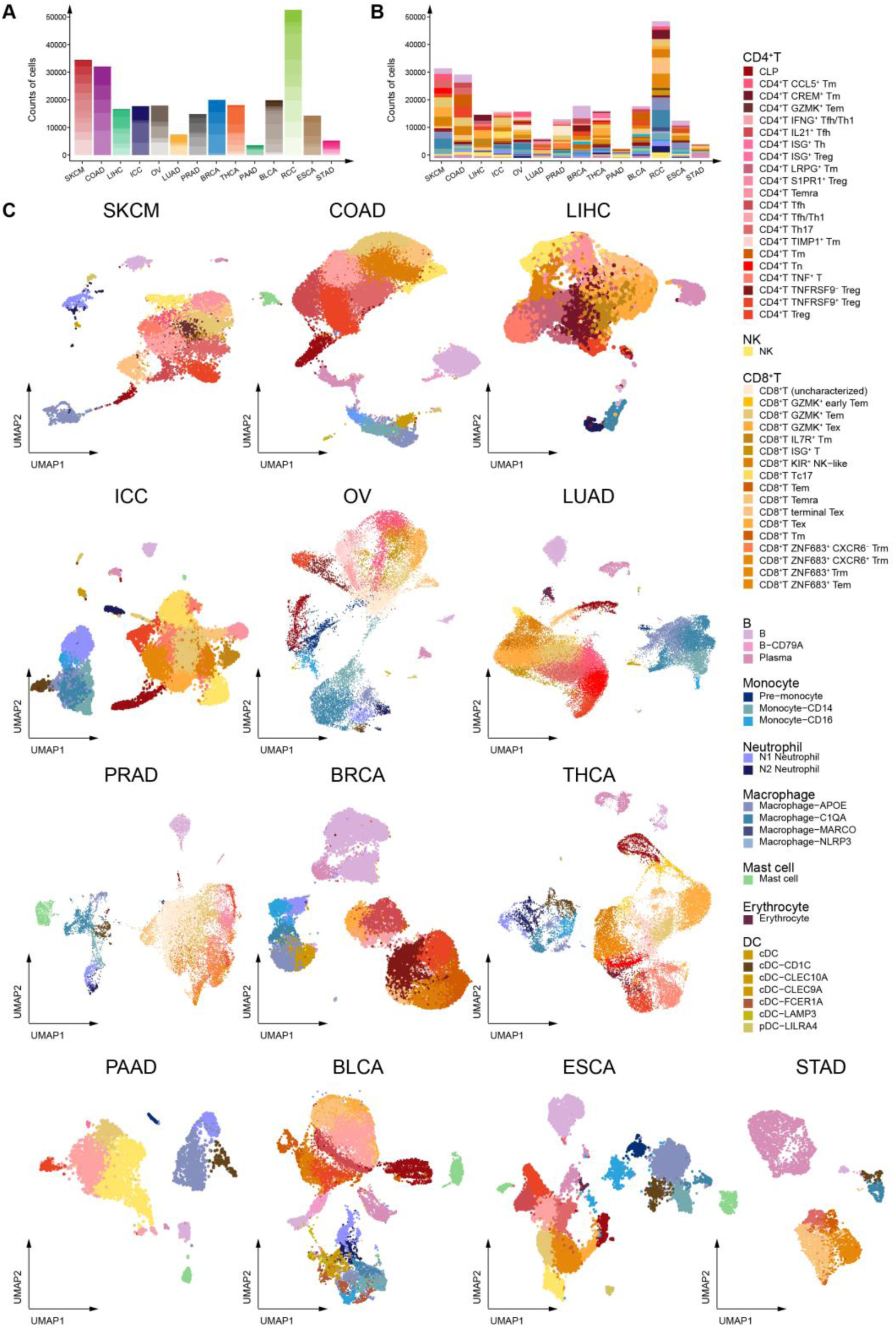
Basic information of the data and hematopoietic/immune cell subsets in each cancer type. (A) Bar plot showing the number of cells collected from different samples in each cancer type. Different levels of transparency indicate different samples in various cancer types. (B) Bar plot showing the number of hematopoietic/immune cells across the cancer types. Similar cell types are represented by the same color scheme. (C) UMAP plots showing the hematopoietic/immune cells in the 13 cancer types.

**Fig. S6.**
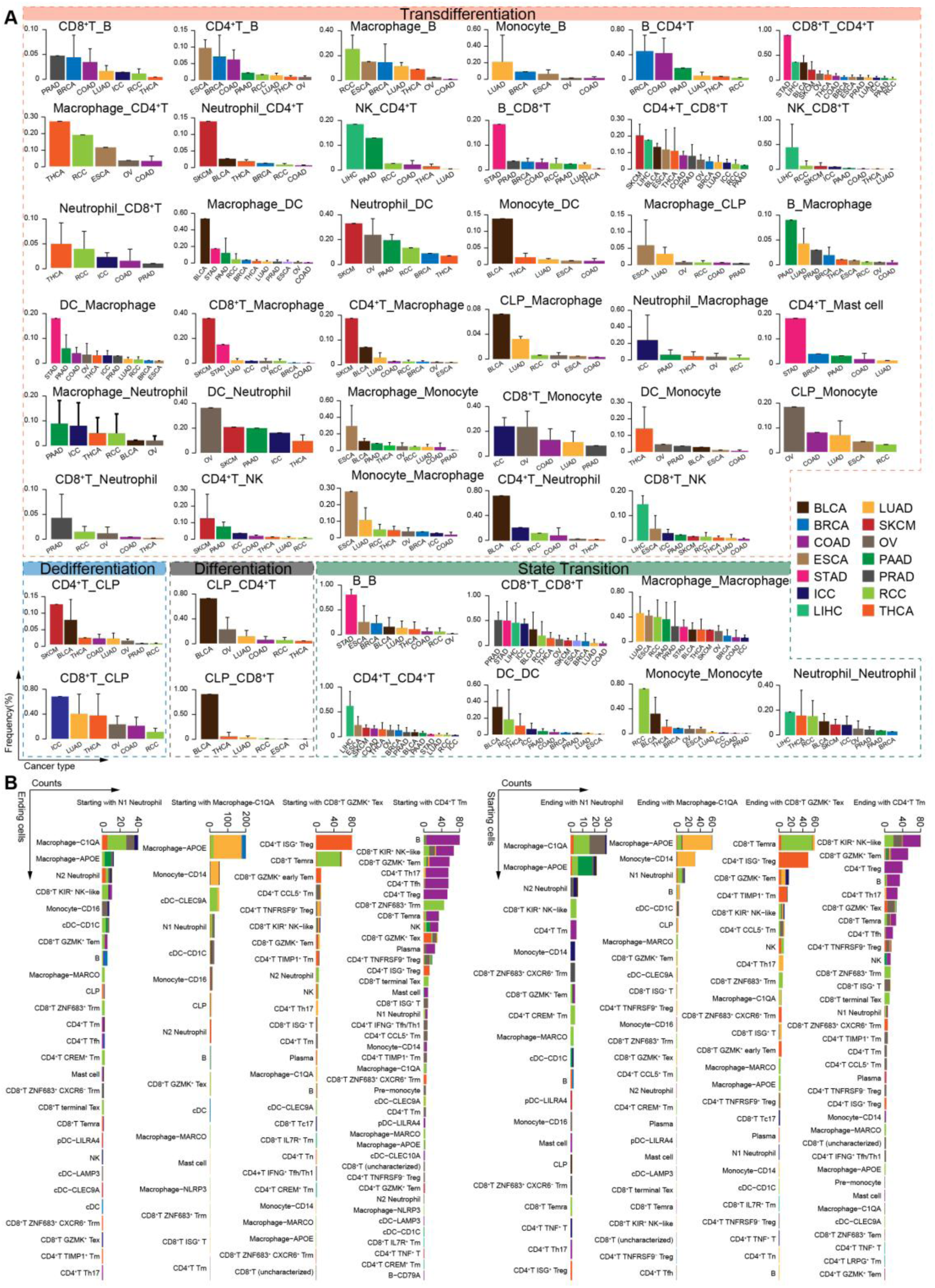
Characteristics and tissue distribution pattern of the cell change types. (A) Bar plots showing the frequency of cell change types in the major lineages across the various cancer types. The cell change types of differentiation, dedifferentiation, state transition and transdifferentiation are highlighted by dotted lines respectively. The color of the dotted lines is the same as the color of the arrows in Fig. 3. The upper side of the error bar is labeled. (B) Bar plots (left) reporting the counts of the ending cells changed from the labelled cell types across various cancer types. Bar plots (right) reporting the counts of the starting cells changed to the labelled cell types across various cancer types.

**Fig. S7.**
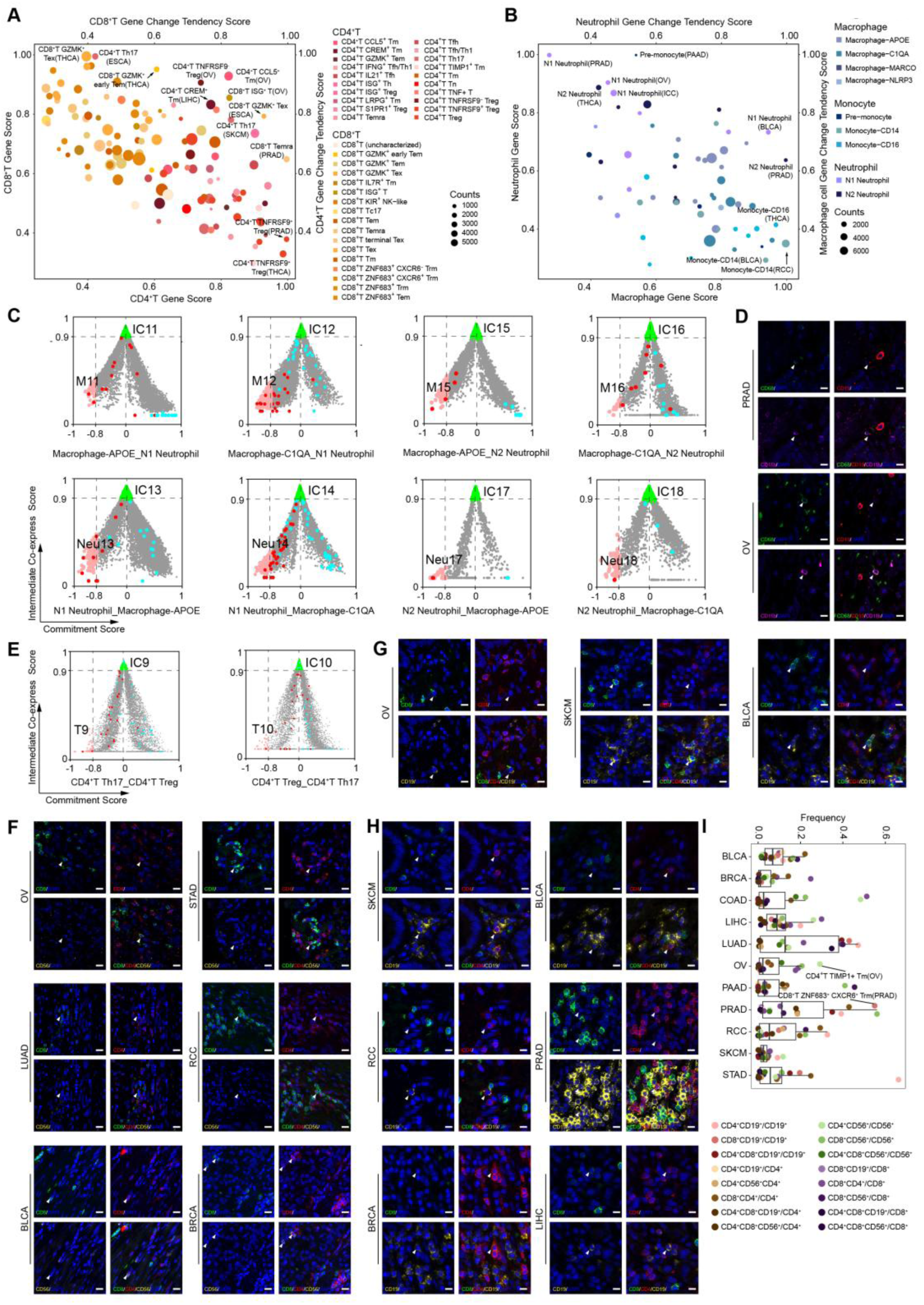
Lymphoid cells and Myeloid cells potential tendency to an intermediate Co-Expressor state. (A) Scatter plot illustrating the gene expression of lymphoid cells and their potential ability to change. Each dot represents one lymphoid cell type of the 14 cancer types and its size is proportional to the number of cells in that particular cell type. For CD4^+^ T cell types, the x-axis represents CD4^+^ T gene score, indicating their gene expression, and the y-axis represents CD4^+^ T gene change tendency score, indicating whether these cells tend to change. For CD8^+^ T cell types, the y-axis represents CD8^+^ T gene score, indicating their gene expression, and the x-axis represents CD8^+^ T gene change tendency score, indicating whether these cells tend to change. (B) Scatter plot displaying the gene expression of myeloid cells and their potential to change. Each dot represents one myeloid cell type of 14 cancer types and the size of the dot corresponds to the number of cells in the cell type. For macrophage and monocyte cell types, the x-axis represents the macrophage gene score, which indicates macrophage gene expression, and the y-axis represents the macrophage gene change tendency score, which indicates whether these cells tend to change. For neutrophil cell types, the y-axis represents the neutrophil gene score, which indicates neutrophil gene expression, and the x-axis represents the neutrophil gene change tendency score, which indicates whether these cells tend to change. (C) Scatter plot showing the transdifferentiation process between macrophage subtypes and neutrophil subtypes. The x-axis represents macrophage/neutrophil cell commitment score, and the y-axis represents the intermediate co-express score. The cells with an x-value less than -0.8 are considered potential starting cells, while the cells with a y-value greater than 0.9 are considered intermediate cells. Green dots represent intermediate cells and pink dots represent potential starting cells. Red dots and blue dots represent the starting and ending cells of the transdifferentiation which we discovered in pan-cancer using the method we built in this study, respectively. (D) Representative examples stained by multiplex immunofluorescence to show the co-expression of CD11b and CD68 (white arrows). The scale bar represents 20 mm. (E) Scatter plot showing the change process of state transition between CD4^+^ T Th17 and CD4^+^ T Treg. The abscissa represents CD4^+^ T Th17/CD4^+^ T Treg cell commitment score, and the ordinate represents Intermediate Co-express Score. Cells with an x-value less than -0.8 are defined as potential starting cells, and cells with a y-value greater than 0.9 are defined as intermediate cells. Green dots represent intermediate cells, pink dots represent potential starting cells, red dots represent the starting cells of the state transition, and blue dots represent the ending cells of the state transition. (F) Representative examples stained by multiplex immunofluorescence to show the co-expression of CD4 and CD8 (white arrows). The scale bar represents 20 mm. (G) Representative examples stained by multiplex immunofluorescence to show the co-expression of CD19 and CD8 (white arrows). The scale bar represents 20 mm. (H) Representative examples stained by multiplex immunofluorescence to show the co-expression of CD19 and CD4 (white arrows). The scale bar represents 20 mm. (I) Box plots showing the proportions of double-positive cells across various cancer types.

**Fig. S8.**
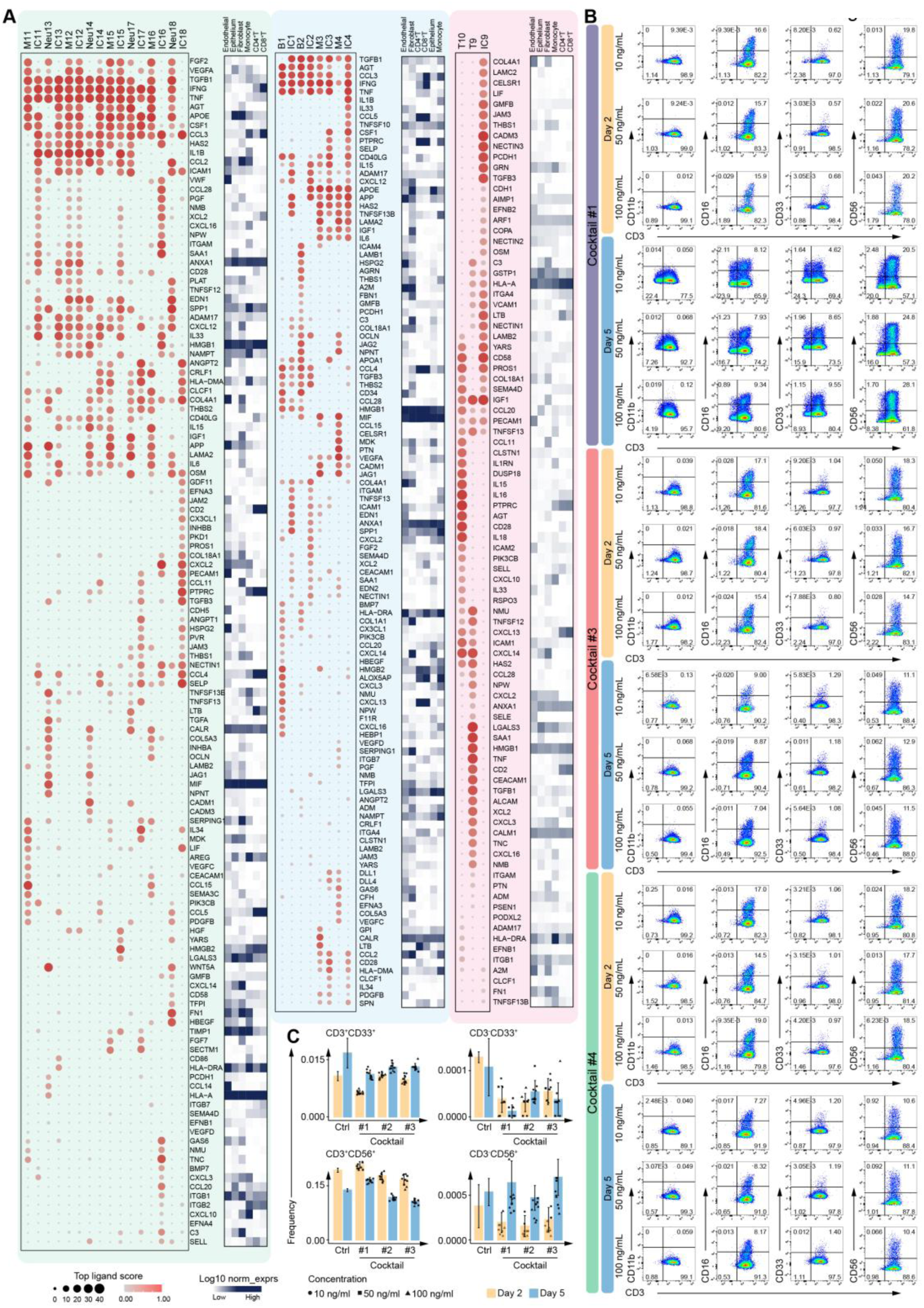
Potential Factors Related to the Cell Plasticity. (A) Bubble plot showing the possible ligands driving the potential starting cells and the intermediate cells against expansion macrophage cells and expansion neutrophil cells, respectively, sent by other immune cells and parenchymal cells (left). Bubble plot illustrating the possible ligands driving the potential starting cells and intermediate cells against expanding B cells and expanding macrophages, respectively, sent by other immune cells and non-hematopoietic cells (middle). Bubble plot illustrating the possible ligands driving the potential starting cells and intermediate cells against expansion CD4^+^ T Th17 cells and expansion CD4^+^ T Treg cells respectively, sent by other immune cells and parenchymal cells (right). The top 40 ligands are selected for each cell type and the heatmap shows these ligands’ expression in the immune cells and parenchymal cells. (B) Representative flow cytometry plots showing phenotype of T cells treated with cocktail #1, #3 and #4 at different concentrations on day 2 and day 5. (C) Bar plots illustrating percentages of CD3^-^ CD56^+^, CD3^-^ CD33^+^, CD3^+^ CD33^+^ and CD3^+^ CD56^+^ cells with cocktail #1, #3 and #4 at different concentrations on day 2 and day 5.

**Fig. S9.**
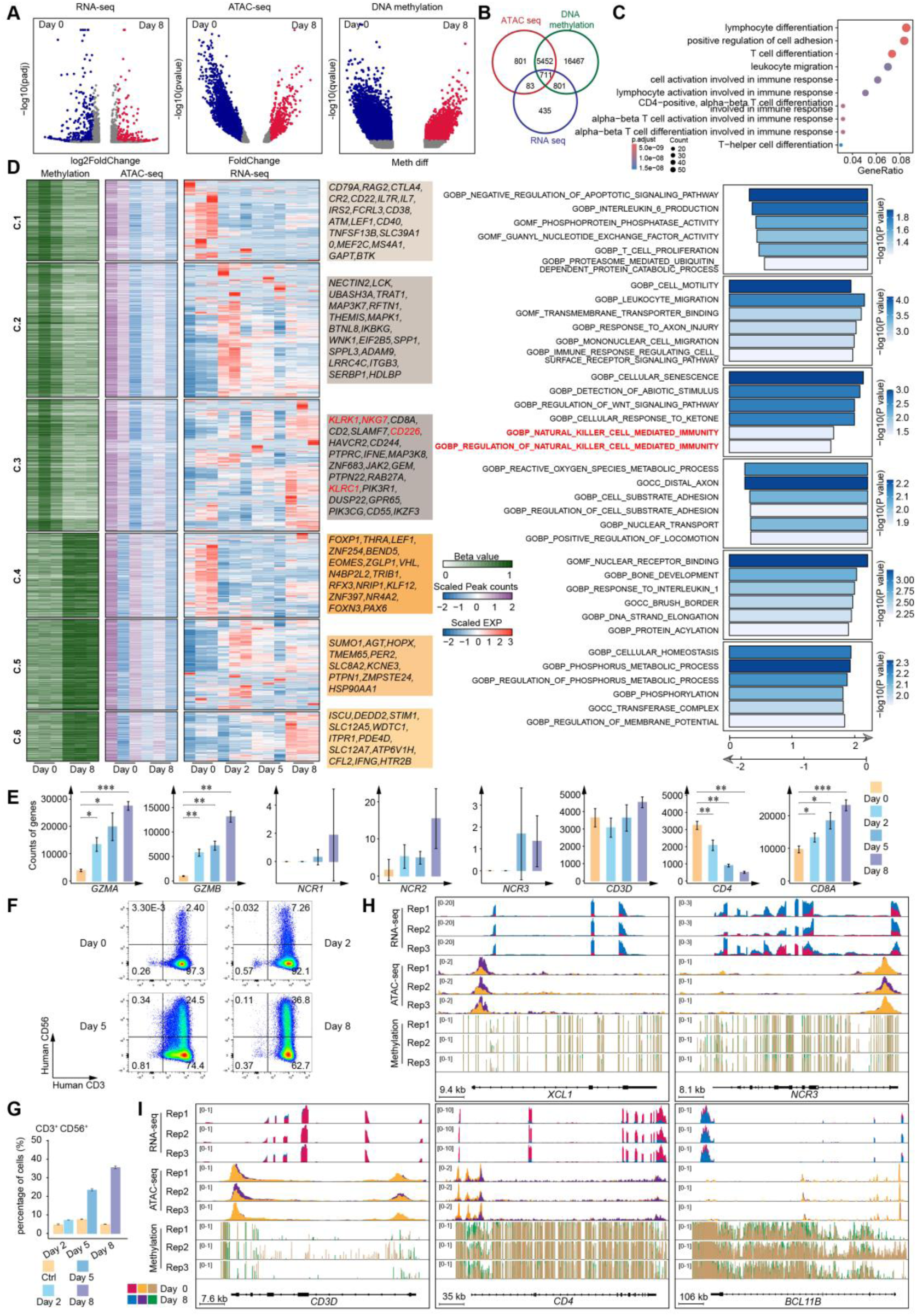
Cocktail-Induced Alterations in DNA Methylation and in T and NK Cell-Associated Genes. (A) Volcano plots illustrating the differentially expressed genes (DEGs), differentially accessible regions (DARs) and differentially methylated positions (DMPs) between day 0 sample and day 8 sample using Benjamini-Hochberg adjusted two-sided Wilcoxon test. (B) Venn diagram showing the overlap of DEGs, DARs and DMPs in (A). (C) GO analysis showing the function of the 711 overlapped genes in (B). (D) Heatmap showing the methylation levels of promoters containing DMPs, defined by the six clusters from K-means clustering analysis. Heatmap of gene expression levels (RNA-seq) and chromatin accessibility (ATAC-seq) were shown according the six clusters. Representative genes were labeled in each cluster. The beta values, scaled gene expression and scaled peak counts were shown. GSEA analysis were shown based each cluster. The pathways associated with NK cells were highlighted. (E) Bar plots showing the cytotoxic factors (*GZMA* and *GZMB*) significantly increased over time. Bar plots showing the NK cell-related genes (*NCR1*, *NCR2* and *NCR3*) slightly increased over time. Bar plots showing the T cell-related genes (*CD3D*, *CD4* and *CD8A*) changed over time. (F) Representative flow cytometry plots showing CD3^+^CD56^+^ cells (NK-like cells) increased with cocktail #2 treatment over time. (G) Bar plots showing the percentages of CD3^+^CD56^+^ cells (NK-like cells) significantly increased over time. (H) Transcriptome (RNA-seq), chromatin accessibility (ATAC-seq) and DNA methylation (WGBS) profiles at the NK cell-related genes *XCL1* and *NCR3* locus in T cells after the cocktail #2 treatment. (I) Transcriptome (RNA-seq), chromatin accessibility (ATAC-seq) and DNA methylation (WGBS) profiles at the T cell-related genes *CD3D*, *CD4* and *BCL11B* locus in T cells after the cocktail #2 treatment.

## References

1. Ren X, Zhang L, Zhang Y, Li Z, Siemers N, Zhang Z: Insights Gained from Single-Cell Analysis of Immune Cells in the Tumor Microenvironment. Annual review of immunology 2021, 39:583–609.

2. Nguyen PHD, Ma S, Phua CZJ, Kaya NA, Lai HLH, Lim CJ, Lim JQ, Wasser M, Lai L, Tam WL et al: Intratumoural immune heterogeneity as a hallmark of tumour evolution and progression in hepatocellular carcinoma. Nature communications 2021, 12(1):227.

3. Cassetta L, Pollard JW: A timeline of tumour-associated macrophage biology. Nature reviews Cancer 2023, 23(4):238–257.

4. Hedrick CC, Malanchi I: Neutrophils in cancer: heterogeneous and multifaceted. Nature reviews Immunology 2022, 22(3):173–187.

5. Xiong S, Dong L, Cheng L: Neutrophils in cancer carcinogenesis and metastasis. Journal of hematology & oncology 2021, 14(1):173.

6. Weinreb C, Rodriguez-Fraticelli A, Camargo FD, Klein AM: Lineage tracing on transcriptional landscapes links state to fate during differentiation. Science (New York, NY) 2020, 367(6479).

7. Sánchez Alvarado A, Yamanaka S: Rethinking differentiation: stem cells, regeneration, and plasticity. Cell 2014, 157(1):110–119.

8. Riddell J, Gazit R, Garrison BS, Guo G, Saadatpour A, Mandal PK, Ebina W, Volchkov P, Yuan GC, Orkin SH et al: Reprogramming committed murine blood cells to induced hematopoietic stem cells with defined factors. Cell 2014, 157(3):549–564.

9. Zhang M, Dong Y, Hu F, Yang D, Zhao Q, Lv C, Wang Y, Xia C, Weng Q, Liu X et al: Transcription factor Hoxb5 reprograms B cells into functional T lymphocytes. Nature immunology 2018, 19(3):279–290.

10. Xie H, Ye M, Feng R, Graf T: Stepwise reprogramming of B cells into macrophages. Cell 2004, 117(5):663–676.

11. Zhou Y, Zhu X, Dai Y, Xiong S, Wei C, Yu P, Tang Y, Wu L, Li J, Liu D et al: Chemical Cocktail Induces Hematopoietic Reprogramming and Expands Hematopoietic Stem/Progenitor Cells. *Advanced science (Weinheim, Baden-Wurttemberg*, Germany) 2020, 7(1):1901785.

12. Qin J, Zhang J, Jiang J, Zhang B, Li J, Lin X, Wang S, Zhu M, Fan Z, Lv Y et al: Direct chemical reprogramming of human cord blood erythroblasts to induced megakaryocytes that produce platelets. Cell stem cell 2022, 29(8):1229–1245.e1227.

13. Wei C, Yu P, Cheng L: Hematopoietic Reprogramming Entangles with Hematopoiesis. Trends in cell biology 2020, 30(10):752–763.

14. Long H, Jia Q, Wang L, Fang W, Wang Z, Jiang T, Zhou F, Jin Z, Huang J, Zhou L et al: Tumor-induced erythroid precursor-differentiated myeloid cells mediate immunosuppression and curtail anti-PD-1/PD-L1 treatment efficacy. Cancer cell 2022, 40(6):674–693.e677.

15. Park YJ, Song B, Kim YS, Kim EK, Lee JM, Lee GE, Kim JO, Kim YJ, Chang WS, Kang CY: Tumor microenvironmental conversion of natural killer cells into myeloid-derived suppressor cells. Cancer research 2013, 73(18):5669–5681.

16. Chen C, Park B, Ragonnaud E, Bodogai M, Wang X, Zong L, Lee JM, Beerman I, Biragyn A: Cancer co-opts differentiation of B-cell precursors into macrophage-like cells. Nature communications 2022, 13(1):5376.

17. Feng Y, Liu G, Li H, Cheng L: The landscape of cell lineage tracing. Science China Life Sciences 2025:1–23.

18. Cheng S, Li Z, Gao R, Xing B, Gao Y, Yang Y, Qin S, Zhang L, Ouyang H, Du P et al: A pan-cancer single-cell transcriptional atlas of tumor infiltrating myeloid cells. Cell 2021, 184(3):792–809.e723.

19. Kourtis N, Wang Q, Wang B, Oswald E, Adler C, Cherravuru S, Malahias E, Zhang L, Golubov J, Wei Q et al: A single-cell map of dynamic chromatin landscapes of immune cells in renal cell carcinoma. Nature cancer 2022, 3(7):885–898.

20. Zheng L, Qin S, Si W, Wang A, Xing B, Gao R, Ren X, Wang L, Wu X, Zhang J et al: Pan-cancer single-cell landscape of tumor-infiltrating T cells. Science (New York, NY) 2021, 374(6574):abe6474.

21. Cliff ERS, Kelkar AH, Russler-Germain DA, Tessema FA, Raymakers AJN, Feldman WB, Kesselheim AS: High Cost of Chimeric Antigen Receptor T-Cells: Challenges and Solutions. Am Soc Clin Oncol Educ Book 2023, 43:e397912.

22. Ludwig LS, Lareau CA, Ulirsch JC, Christian E, Muus C, Li LH, Pelka K, Ge W, Oren Y, Brack A et al: Lineage Tracing in Humans Enabled by Mitochondrial Mutations and Single-Cell Genomics. Cell 2019, 176(6):1325–1339.e1322.

23. Xu J, Nuno K, Litzenburger UM, Qi Y, Corces MR, Majeti R, Chang HY: Single-cell lineage tracing by endogenous mutations enriched in transposase accessible mitochondrial DNA. eLife 2019, 8.

24. Magen A, Hamon P, Fiaschi N, Soong BY, Park MD, Mattiuz R, Humblin E, Troncoso L, D’souza D, Dawson T: Intratumoral dendritic cell–CD4+ T helper cell niches enable CD8+ T cell differentiation following PD-1 blockade in hepatocellular carcinoma. Nature Medicine 2023:1–11.

25. Zhao T, Fu Y, Zhu J, Liu Y, Zhang Q, Yi Z, Chen S, Jiao Z, Xu X, Xu J et al: Single-Cell RNA-Seq Reveals Dynamic Early Embryonic-like Programs during Chemical Reprogramming. Cell stem cell 2018, 23(1):31–45.e37.

26. Raghavan S, Winter PS, Navia AW, Williams HL, DenAdel A, Lowder KE, Galvez-Reyes J, Kalekar RL, Mulugeta N, Kapner KS et al: Microenvironment drives cell state, plasticity, and drug response in pancreatic cancer. Cell 2021, 184(25):6119–6137.e6126.

27. Bendelac A, Savage PB, Teyton L: The biology of NKT cells. Annual review of immunology 2007, 25:297–336.

28. Gagliani N, Amezcua Vesely MC, Iseppon A, Brockmann L, Xu H, Palm NW, de Zoete MR, Licona-Limón P, Paiva RS, Ching T et al: Th17 cells transdifferentiate into regulatory T cells during resolution of inflammation. Nature 2015, 523(7559):221–225.

29. Flavell RA, Sanjabi S, Wrzesinski SH, Licona-Limón P: The polarization of immune cells in the tumour environment by TGFbeta. Nature reviews Immunology 2010, 10(8):554–567.

30. De Carvalho DD, You JS, Jones PA: DNA methylation and cellular reprogramming. Trends in cell biology 2010, 20(10):609–617.

31. Li Y, Wang J, Zhou L, Gu W, Qin L, Peng D, Li S, Zheng D, Wu Q, Long Y: DNMT1 inhibition reprograms T cells to NK-like cells with potent antitumor activity. Science Immunology 2025, 10(105):eadm8251.

32. Griffin GK, Booth CA, Togami K, Chung SS, Ssozi D, Verga JA, Bouyssou JM, Lee YS, Shanmugam V, Hornick JL: Ultraviolet radiation shapes dendritic cell leukaemia transformation in the skin. Nature 2023:1–8.

33. Rao S, Lee SY, Gutierrez A, Perrigoue J, Thapa RJ, Tu Z, Jeffers JR, Rhodes M, Anderson S, Oravecz T et al: Inactivation of ribosomal protein L22 promotes transformation by induction of the stemness factor, Lin28B. Blood 2012, 120(18):3764–3773.

34. Dong L, Wei C, Xiong S, Yu P, Zhou R, Cheng L: Spliceosome inhibitor induces human hematopoietic progenitor cell reprogramming toward stemness. Exp Hematol Oncol 2022, 11(1):37.

35. Zhou Y, Wei C, Xiong S, Dong L, Chen Z, Chen SJ, Cheng L: Trajectory of chemical cocktail-induced neutrophil reprogramming. Journal of hematology & oncology 2020, 13(1):171.

36. Yoshida Y, Takahashi K, Okita K, Ichisaka T, Yamanaka S: Hypoxia enhances the generation of induced pluripotent stem cells. Cell stem cell 2009, 5(3):237–241.

37. Zhang X, Cao D, Xu L, Xu Y, Gao Z, Pan Y, Jiang M, Wei Y, Wang L, Liao Y et al: Harnessing matrix stiffness to engineer a bone marrow niche for hematopoietic stem cell rejuvenation. Cell stem cell 2023, 30(4):378–395.e378.

38. Tang Y, Xiong S, Yu P, Liu F, Cheng L: Direct Conversion of Mouse Fibroblasts into Neural Stem Cells by Chemical Cocktail Requires Stepwise Activation of Growth Factors and Nup210. Cell reports 2018, 24(5):1355–1362.e1353.

39. Xiong S, Feng Y, Cheng L: Cellular Reprogramming as a Therapeutic Target in Cancer. Trends in cell biology 2019, 29(8):623–634.

40. Yang D, Jones MG, Naranjo S, Rideout WM, 3rd, Min KHJ, Ho R, Wu W, Replogle JM, Page JL, Quinn JJ et al: Lineage tracing reveals the phylodynamics, plasticity, and paths of tumor evolution. Cell 2022, 185(11):1905–1923.e1925.

41. Macaulay IC, Haerty W, Kumar P, Li YI, Hu TX, Teng MJ, Goolam M, Saurat N, Coupland P, Shirley LM et al: G&T-seq: parallel sequencing of single-cell genomes and transcriptomes. Nature methods 2015, 12(6):519–522.

42. Wang K, Ye R, Bai S, Xiao Z, Yang L, Li J, Tang C, Sei E, Peng J, Casasent AK et al: Coalescing single-cell genomes and transcriptomes to decode breast cancer progression. Cell 2025, 188(22):6355–6369.e6316.

43. Kester L, van Oudenaarden A: Single-Cell Transcriptomics Meets Lineage Tracing. Cell stem cell 2018, 23(2):166–179.

